# Increasing metadata coverage of SRA BioSample entries using deep learning based Named Entity Recognition

**DOI:** 10.1101/414136

**Authors:** Adam Klie, Brian Y Tsui, Shamim Mollah, Dylan Skola, Michelle Dow, Chun-Nan Hsu, Hannah Carter

**Affiliations:** Department of Medicine, Division of Medical Genetics, University of California San Diego, La Jolla, CA 92093; Bioinformatics and Systems Biology Program, University of California San Diego, La Jolla, CA 92093; Department of Bioengineering, University of California San Diego, La Jolla, CA 92093; Department of Genetics, Washington University in St. Louis, St. Louis, MO 63130; Department of Neurosciences, University of California San Diego, La Jolla, CA 92093

## Abstract

High quality metadata annotations for data hosted in large public repositories are essential for research reproducibility, and for conducting fast, powerful and scalable meta-analyses. Currently, a majority of sequencing samples in the National Center for Biotechnology Information’s (NCBI’s) Sequence Read Archive (SRA) are missing metadata across several categories. In an effort to improve the metadata coverage of these samples, we leveraged almost 44 million attribute-value pairs from SRA BioSample to train a scalable, recurrent neural network that predicts missing metadata via Named Entity Recognition (NER). The network was first trained to classify short text phrases according to 11 metadata categories and achieved an overall accuracy and area under the receiver operating characteristic (AUROC) curve of 85.2% and 0.977 respectively. We then applied our classifier to predict 11 metadata categories from the longer TITLE attribute of samples, evaluating performance on a set of samples withheld from model training. Prediction accuracies were high when extracting sample Species (94.85%), Condition/Disease (95.65%) and Strain (82.03%) from TITLEs, with lower accuracies and lack of predictions for other categories highlighting multiple issues with the current metadata annotations in BioSample. These results indicate the utility of recurrent neural networks for NER-based metadata prediction and the potential for models such as the one presented here to increase metadata coverage in BioSample while minimizing the need for manual curation.

**Availability:** All the analyses, environments, and Jupyter notebooks pertaining to this manuscript are available on Github: https://github.com/cartercompbio/PredictMEE.

## Introduction

Advances in next-generation sequencing (NGS) technologies have led to the rapid accumulation of publicly available sequencing datasets. Large repositories, such as the National Center for Biotechnology Information’s (NCBI) Gene Expression Omnibus (GEO) (1) or Sequence Read Archive (SRA) (2), have been established to store the raw and processed versions of this data, allowing researchers to easily reanalyze and repurpose existing datasets. High quality metadata associated with these datasets is essential for this repurposing, allowing for greater reproducibility, for targeted studies of specific phenotypes, and for large-scale meta-analysis across studies. Paired with efforts to uniformly normalize the preprocessing of raw sequencing data (3, 4), comprehensive and accurate metadata has the potential to open the door for many powerful and targeted analyses to address important biological questions and problems.

Ideally, metadata necessary to accurately describe the technical and biological variation in a given sample would be available for every sample in these repositories. However, the current state of metadata quality in many archived samples fails to meet this standard. Gonçalves *et al*. (5) recently described the variable state of the metadata available in databases such as NCBI’s BioSample and the European Bioinformatics Institute’s (EBI) BioSamples (6). The infrequent use of controlled vocabularies in the metadata submission process, coupled by the allowance for the creation of user-defined attributes, has led to an explosion of heterogeneity in the overall metadata landscape (5). This can often hinder researchers’ ability to fully utilize the potential information that a given dataset, or a meta-analysis of multiple datasets, might hold.

This database heterogeneity has inspired several recent efforts to improve metadata of future stored datasets by remodeling the current submission process. Various biological sub-disciplines have established guidelines to help standardize the ontologies researchers use in reporting relevant metadata upon submission (7, 8). Bukhari *et al.* (9) have also developed a web-based browser plug-in that recommends metadata to the user, where recommendations are native to a given repository and are based on sets of standard ontologies. For NCBI, the BioSample (10) database was established to promote the standardization of annotation used to characterize the data for samples deposited in GEO, SRA and other archives hosted by NCBI.

Strategies to remedy the current metadata landscape of these repositories are also actively being pursued. These strategies can typically be binned into one of three categories: [1] manual curation, [2] automated or semi-automated curation, or [3] inferring metadata from the underlying raw sample data (often gene expression data). Manual curation remains the most accurate solution (11), but does not scale with current data quantities. Many automated or semi-automated methods attempt to normalize the metadata by clustering or mapping to ontologies (12, 13) and scale much better with increasing data quantities. Methods for increasing completeness using automated or semi-automated techniques often center on utilizing named entity recognition (NER), a Natural Language Processing (NLP) technique used to identify predefined entities in unstructured text, to retrieve metadata entities from the unstructured text associated with a sample. In most cases, however, the aforementioned strategies still require some level of manual annotation, and do not scale to encompass the entirety of the metadata associated with these repositories.

Here, we first analyzed the landscape of SRA metadata in BioSample, which is organized as structured “attribute-value” pairs (e.g., tissue-liver). We found that most attributes had entirely missing values in a large proportion of samples, with substantial heterogeneity noted in both attribute definitions and in the values of metadata within a given attribute. Due to the availability of 43,907,007 attribute-value pairs and the power of neural networks in automating NLP tasks, we opted to use deep learning to locate and extract a set of relevant metadata categories, such as species or sex, from longer free-text attributes, such as sample titles and descriptions. First, we trained a recurrent neural network to classify short phrases into 11 metadata categories and achieved an accuracy and area under the receiver operating characteristic (AUROC) curve of 85.2% and 0.977 respectively. We then used the trained classifier to perform NER on the longer TITLE attributes associated with each sample and found that we could achieve high accuracy metadata prediction of Species (94.85%), Condition/Disease (95.65%) and Strain (82.03%). Lower predictive performance for another eight metadata categories proved to be highly illustrative of issues with current metadata annotations in BioSample SRA (Table 1). Our results illustrate current limitations in coverage and consistency of BioSample metadata in SRA and indicate that a deep neural network trained on a large dataset has the potential to significantly increase metadata coverage in SRA with minimal manual curation.

**Table 1.**
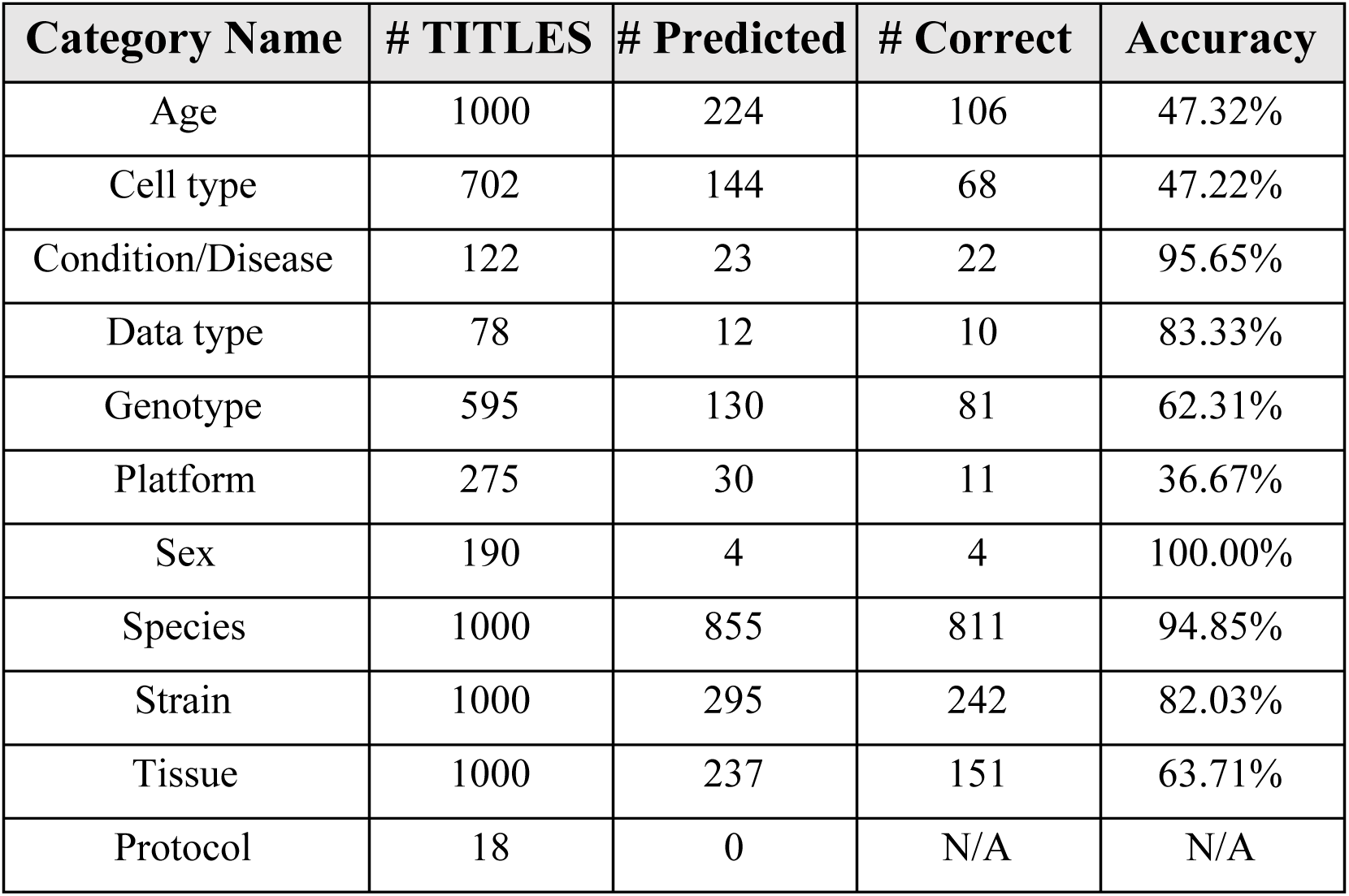
Performance on the prediction of the 11 metadata categories from TITLEs

| Category Name | # TITLES | # Predicted | # Correct | Accuracy |
| --- | --- | --- | --- | --- |
| Age | 1000 | 224 | 106 | 47.32% |
| Cell type | 702 | 144 | 68 | 47.22% |
| Condition/Disease | 122 | 23 | 22 | 95.65% |
| Data type | 78 | 12 | 10 | 83.33% |
| Genotype | 595 | 130 | 81 | 62.31% |
| Platform | 275 | 30 | 11 | 36.67% |
| Sex | 190 | 4 | 4 | 100.00% |
| Species | 1000 | 855 | 811 | 94.85% |
| Strain | 1000 | 295 | 242 | 82.03% |
| Tissue | 1000 | 237 | 151 | 63.71% |
| Protocol | 18 | 0 | N/A | N/A |

## Materials and Methods

### BioSample Data

Each BioSample SRA entry is a record of metadata associated with a single biospecimen in SRA. Metadata associated with a specimen are broken up into attribute-value pairs, where the attribute specifies the metadata type (age, cell type, sex, etc.) and the values are the corresponding metadata associated with that type (Figure 1A). As of May 15, 2018, 43,907,007 such attribute-value pairs were available for download from NCBI (https://ftp.ncbi.nlm.nih.gov/sra/reports/Metadata/), encompassing 2,912,000 samples and over 100,000 studies.

**Figure 1.**
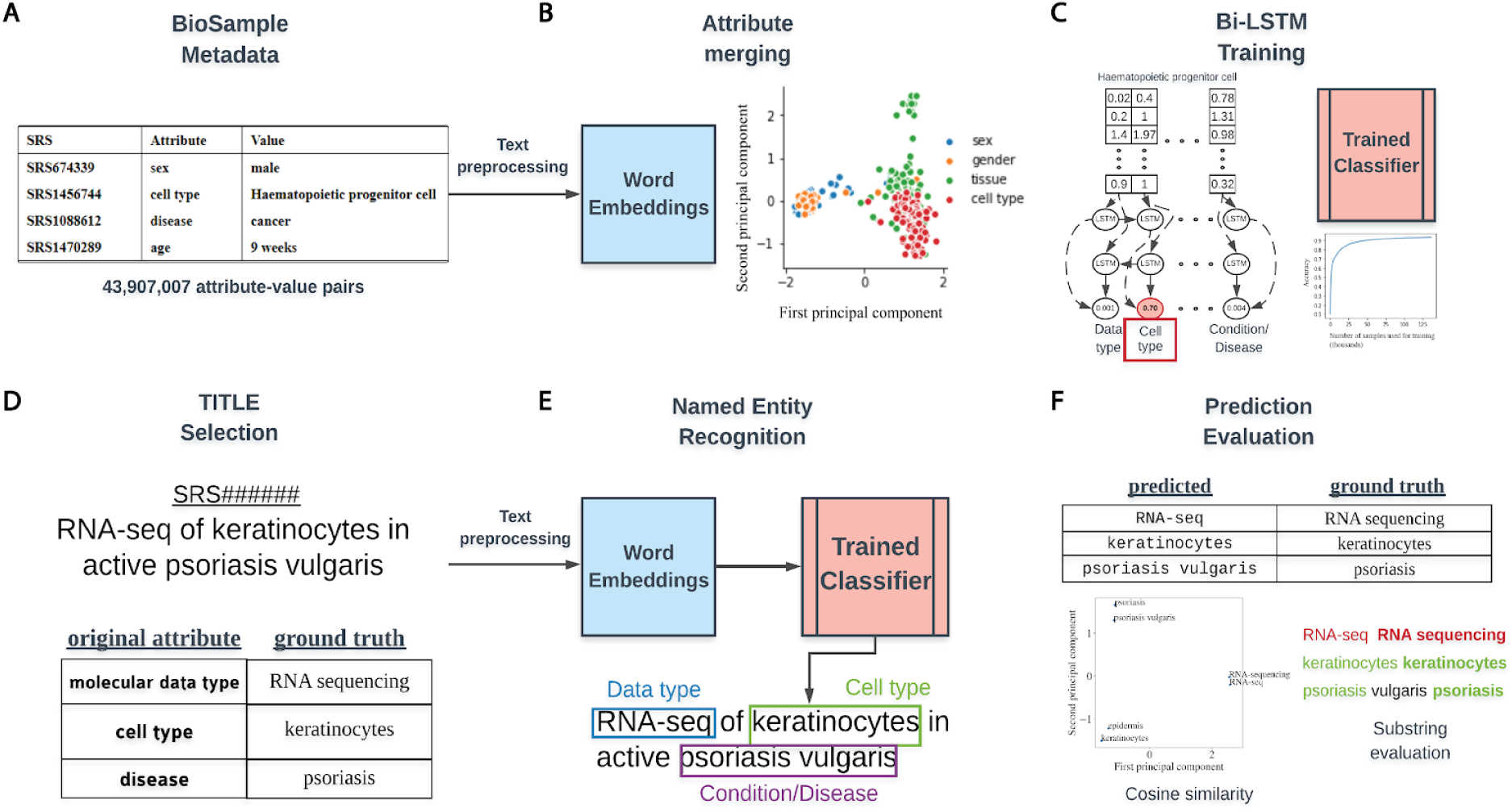
Overview of classifier training and metadata prediction workflow. **(A)** A few examples of the 44 million attribute-value pairs in SRA BioSample. **(B)** Word embeddings of preprocessed values allowed for the clustering and merging of attributes that were similar in the embedding space. **(C)** A subset of attribute-value pairs was split into a training and test set and a bi-LSTM classifier was trained to identify 11 metadata categories. **(D)** TITLEs were selected as the free text for NER using the trained model. An example TITLE with associated ground truth labels is shown. **(E)** These TITLEs were preprocessed into n-grams and fed into the trained classifier after word embedding to generate metadata predictions for the 11 categories. **(F)** Comparisons to ground truth metadata were done using substring matching and cosine similarity in the word embedding space.

The NCBI defines 456 attributes (https://www.ncbi.nlm.nih.gov/biosample/docs/attributes/) for common use in annotating samples in SRA, with users defining all others upon submission. We selected 11 attributes as a basis for classes in training a classifier for prediction of metadata from sample TITLEs, which are often brief summaries describing samples. These attributes were age, cell type, disease, molecular data type, genotype, platform, protocol, sex, SCIENTIFIC_NAME, strain and tissue. These attributes are referred to as selected attributes hereafter (Supplementary Table 1, Selected Attribute).

### Word Embeddings and Attribute Merging

Natural language processing (NLP) approaches using neural networks require the conversion of free-text to numerical vectors as input. An often-used approach for this task is to vectorize words using a Word2vec embedding model (14). These word embedding models are trained on large corpora (text) to numerically capture contextual and semantic information of words and represent word similarity as a geometric distance in an n-dimensional space. In this way, words that are semantically similar, such as “sex” and “gender”, would be expected to be close in geometric distance in the embedding space (Figure 1B). This approach has the advantage of not requiring hard-coding of the semantic similarity between words, but does require that we use a context specific model. We utilized a publicly available Word2vec model that was trained on the entire PubMed, PMC and Wikipedia text corpora and included 5,443,656 word vectors, each with 200 features (15). We chose the simpler Word2vec model architecture instead of an attention-based transformer model such as BERT (16) because our task involved classifying short phrases with less long-range contextual information present.

Attribute merging was performed on the basis of similarity of values in the 200-dimensional word embedding space. To group like attributes, 100 randomly selected values from all attributes occurring 100 or more times across the dataset were vectorized and averaged to generate a mean embedding vector representative of each attribute. The cosine similarity between the mean vectors of the 11 selected attributes with the mean vectors for all other attributes was then calculated. All the attributes with a similarity of 0.8 or greater to each selected attribute were merged together to create groupings of attributes representing the same concept. These meta-attribute groupings were each given a label that represents the underlying concept the attributes in the grouping shared. These groupings are hereafter referred to as metadata categories and labeled by their category name (Supplementary Table 1, Category Name).

### Bi-LSTM Training

To classify short text phrases according to the 11 metadata categories, we constructed a bi-directional long-short term memory (bi-LSTM) recurrent neural network and trained it on BioSample values that had a word length constrained between 2 and 7 (Figure 1C). No study was allowed to contribute more than 100 samples to the training set to avoid study bias (Supplementary Figure 1A) and the number of training examples for each metadata category was capped at 20,000 to avoid the model learning to predict categories with more training examples by default (Supplementary Figure 1B). To build and validate the classifier, a 4:1 split by study of examples passing the above criteria was used for the training and test sets respectively.

Training and test set values were first encoded by word embedding IDs. These ID vectors were then fed into the following layers of the bi-LSTM:

1. An embedding layer that converted word embedding IDs to word vectors
2. A bidirectional layer with 64 hidden units and dropout rate of 0.5
3. A dense layer with a logistic activation function to output the probability score of each category

The Adam optimizer with categorical cross-entropy as a loss function was used for training a single epoch with a batch size of 100 and a learning rate of 0.001. Keras (v.2.2.2) with tensorflow (v.1.9.0) backend was used for model construction, training and testing. Evaluation of training performance was done using standard multi-class classification metrics such as precision, recall, F1-score and AUROC. The average AUROC value reported was calculated on an aggregate of all test set examples and took into account the differing number of examples in each class.

### Preprocessing, Metadata Prediction, and Evaluation of Performance

The trained classifier could then be used to locate the 11 metadata categories from longer, unstructured text using NER. We selected values from the TITLE attribute of samples (Figure 1D) as sources for extraction, keeping only TITLEs greater than 5 words in length. Before metadata prediction, TITLEs were preprocessed by first splitting them into sentences based on the common sentence delimiters ‘;’’,’’.’. All whitespaces were then replaced with a single whitespace character, ‘’ and sentences were tokenized using the python nltk package (v3.4.5). Any empty tokens or stop words were also removed.

Prediction of metadata labels from TITLEs was done with an n-gram approach (Figure 1E). For all possible n-grams of length 2 to 7 in each sentence of an input TITLE, we applied the trained model to assign a score to that n-gram for each of the 11 metadata categories. Any prediction scores that were within 0.01 of the prediction scores for the empty string (‘’) were discarded, and only n-grams that had at least 2 tokens within the vocabulary of the Word2vec model were considered. Each n-gram was assigned to the metadata category (e.g., Species, Age, Disease/Condition) with the highest score calculated by the model. To remove low confidence predictions, we removed any n-grams for which the difference in the highest and second highest scoring categories was less than or equal to 0.1. If multiple n-grams were classified to the same metadata category, the highest scoring n-gram of all overlapping n-grams was retained, with all others discarded.

To evaluate the performance of this prediction algorithm, we sampled 1,000 TITLEs for most of the metadata categories that had “ground truth” values already annotated for that category within BioSample. In other words, the sample for each selected TITLE had to have at least one of the selected attributes (e.g., molecular data type) already annotated for the TITLE to be included in the evaluation. The underlying metadata value for this attribute was considered to be the ground truth that we could compare our prediction against. Some selected attributes had far less than 1,000 TITLEs that met these criteria, and for these, as many TITLEs as possible were kept. Evaluation of string similarity remains a challenge in NLP, and due to the various ways in which the metadata is annotated in BioSample SRA, exact matching between predicted and actual values was too stringent of a measure to determine prediction accuracy. Instead, if the actual value had a cosine similarity in the word-embedding space of 0.7 or more to the predicted value, or if the entire predicted value was a substring contained within the ground truth value (or vice-versa), it was considered a match (Figure 1F).

## Results

### A large proportion of metadata in BioSample SRA is missing

Metadata associated with SRA is hosted by NCBI’s BioSample to store annotations of sequencing data. These annotations are represented by attribute-value pairs, where the “attribute” refers to a category of metadata, and “value” refers to the specific annotation under that category describing the underlying sample (Figure 2A). BioSample defines 456 such attribute categories, but also allows submitters the option of supplying their own attributes through their submission portal. We downloaded a snapshot of the SRA metadata stored on the BioSample database from May 15^th^, 2018 and analyzed the landscape of the attribute-value pairs associated with samples. The dataset comprised 43,907,007 attribute-value pairs encompassing 2,921,722 samples in SRA for an average of 15.03 attribute-value pairs per sample. The samples cover 19,361 total unique attributes of which 316 are defined by BioSample. The vast majority of attributes are user-defined (19,045), and only 21.8% of all values in this dataset are paired with BioSample defined attributes. Specifically, the majority of the top 15 most commonly used attributes are user defined (Figure 2B). These data illustrate a growing problem in online repositories such as BioSample, where user-defined fields dominate the metadata landscape.

**Figure 2.**
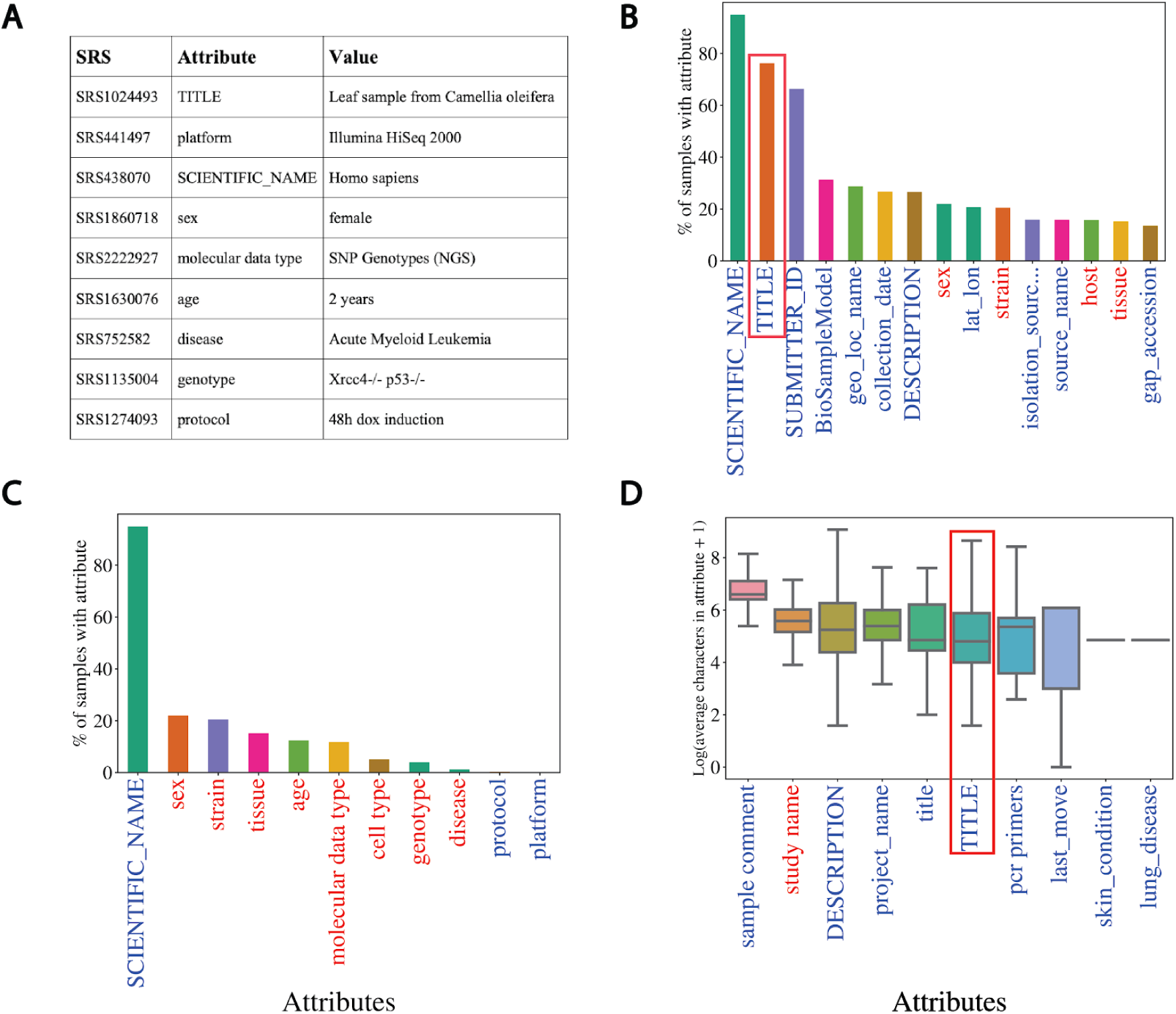
Missing metadata in SRA. **(A)** Examples of SRA attribute-value pairs. Percentage of all samples that contained annotations for the **(B)** top 15 most used attributes and **(C)** the 11 selected attributes. X-axis shows attribute type and y-axis shows the percentage of total samples that used the given attribute. **(D)** Distributions of the average number of characters for the 10 longest (by mean) attributes in BioSample annotations of SRA. X-axis shows attribute type and y-axis shows the Log2(average characters) for a given attribute. Blue labels indicate a user defined attribute, red labels indicate a BioSample defined attribute. TITLE attribute in panels (B) and (D) is highlighted.

We then selected the 11 attributes that we felt best described as much of the biological and technical variation of a sequencing sample for further analysis of metadata coverage (Supplementary Table 1, Selected Attribute). Examining the coverage of these selected attributes across all samples, we see that only “SCIENTIFIC_NAME” (used most frequently to describe a sample’s species) covers more than 25% of samples (Figure 2C). These same trends are observed when we look at only *Homo sapiens* samples (i.e., samples with a SCIENTIFIC_NAME annotated as “Homo sapiens”). In this subset of samples, we see relatively low coverage even in the 15 most used attributes (Supplementary Figure 2A). For the 11 selected attributes, only SCIENTIFIC_NAME and sex are annotated in more than 50% of the *Homo sapien* samples (Supplementary Figure 2B). These observations indicate that missing data is problematic in the BioSample SRA metadata landscape.

Named-entity recognition (NER) presents a potential solution to help remedy the missing metadata problem in SRA BioSample entries. NER requires longer free-text sentences as input to allow for context-specific entities to be detected. The average character length distributions of the 10 longest attributes in BioSample are shown in Figure 2D (Supplementary Figure 2C for *Homo sapien* samples). Given that it is one of the most prevalent and one of the longest attributes in the dataset, the TITLE field represents a potential input for an NER model to extract metadata from. If a sample is missing a given metadata attribute, that annotation may be present and extractable in the TITLE of that sample. Indeed, when we look at the subset of samples of the overall dataset that include a TITLE attribute, we see that almost all of the 11 selected metadata attributes show low coverage (Supplementary Figure 2D). This suggests that the TITLE attribute could be used as a source of missing metadata to increase coverage across SRA BioSample.

### Word embeddings capture the semantic similarity of domain specific words

In order to perform automated NER on sample TITLEs, we utilized a biomedical word embedding model to vectorize text to a numerical input that could be handled by a neural network. We used a Word2vec embedding model from Chiu *et al.* (15) that was pre-trained on PubMed, PMC and Wikipedia, and vectorized words to 200 features. Figure 3 illustrates the word embedding model’s ability to capture the semantic similarity between words relevant to biomedical metadata prediction. Highlighted are the embedding model’s ability to cluster broader categories of entity types such as disease, age, data type and sex (Figure 3A and 3B), as well as more subtle biologically relevant variation in data type (Figure 3C and 3D).

**Figure 3.**
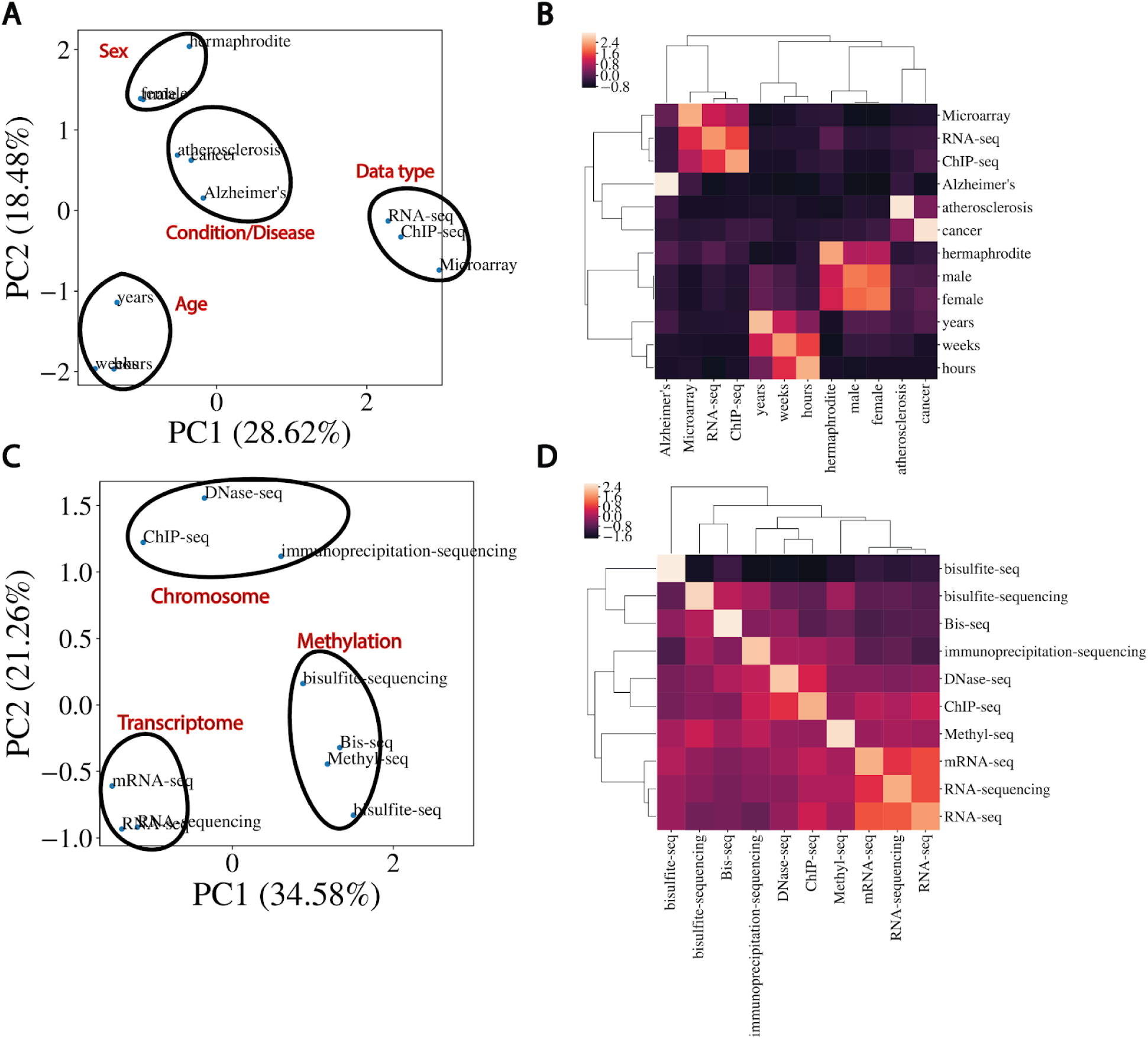
Word embeddings capture semantic similarities between words. **(A)** Dimensional reduction using PCA to visualize vectorized representation of a set of words encapsulating disease, age, data type and sex. **(B)** Corresponding correlation heatmap of the z-scored cosine similarity between words, with hierarchical clustering shown. **(C)** Same as (A) for data types. **(D)** Corresponding correlation heatmap of z-scored cosine similarity between words in (C).

Inconsistencies in naming of attributes that are semantically the same limited the number of examples that could be used for model training for a subset of the selected 11 attributes. These inconsistencies often manifest themselves in differences in capitalization, spelling or punctuation (e.g., cell type and cell_type) and seem to be caused by user-defined attribute submission. We reasoned that the word embedding model could also be utilized to cluster attributes by semantic similarity to increase the pool of training examples for each category. We calculated mean embedding vectors for all attributes followed by the pairwise cosine similarities between them. Merging attributes that had a cosine similarity greater than 0.8 to the 11 selected attributes (Supplementary Table 2) significantly increased the coverage of each selected attribute for model training (Supplementary Table 3). These merged attribute groups were given broader category names describing the semantic concept they represented. These new metadata categories formed the basis of classes for classification output and subsequent prediction of metadata from sample TITLEs.

### A bi-LSTM classifier can classify short text values with relatively high accuracy for most classes

We chose a bi-directional long-short term memory (bi-LSTM) network architecture to classify short text values according to the 11 metadata categories, as bi-LSTMs have been shown to capture the sequential nature of text and the short and long term relationships between words (17). We trained such a classifier to recognize the 11 metadata categories using 133,627 attribute-value pairs and a single epoch, and were able to achieve an overall test set accuracy of 85.2% and an AUROC of 0.977 (Figure 4A and 4B). Greater than 90% classification accuracy was noted for Data type, Age, Species, Sex and Platform, with accuracy in the 80% range observed for Genotype, Strain and Tissue, and sub 70% accuracy observed only for Protocol and Cell type (Figure 4C). Protocol was the class with the fewest training examples, containing only 635, and was often mistaken for Genotype or Tissue. Similarly, the model was unable to consistently distinguish between Cell type and Tissue, predicting Tissue for 22% of Cell type test set examples (Figure 4C). The latter observation is perhaps not surprising given that the cosine similarity between Cell type and Tissue is very high in the word embedding space (Supplementary Figure 3). The merging of semantically similar attributes likely contributes to the high degree of similarity observed between selected attributes. However, high similarities noted prior to attribute merging indicate that this is most likely caused by user difficulty in distinguishing attributes upon submission. Despite these issues, accuracy, precision, recall, F1 score, average AUROC and per-class AUROCs (Figure 4D) all remained high, indicating good classification performance overall.

**Figure 4.**
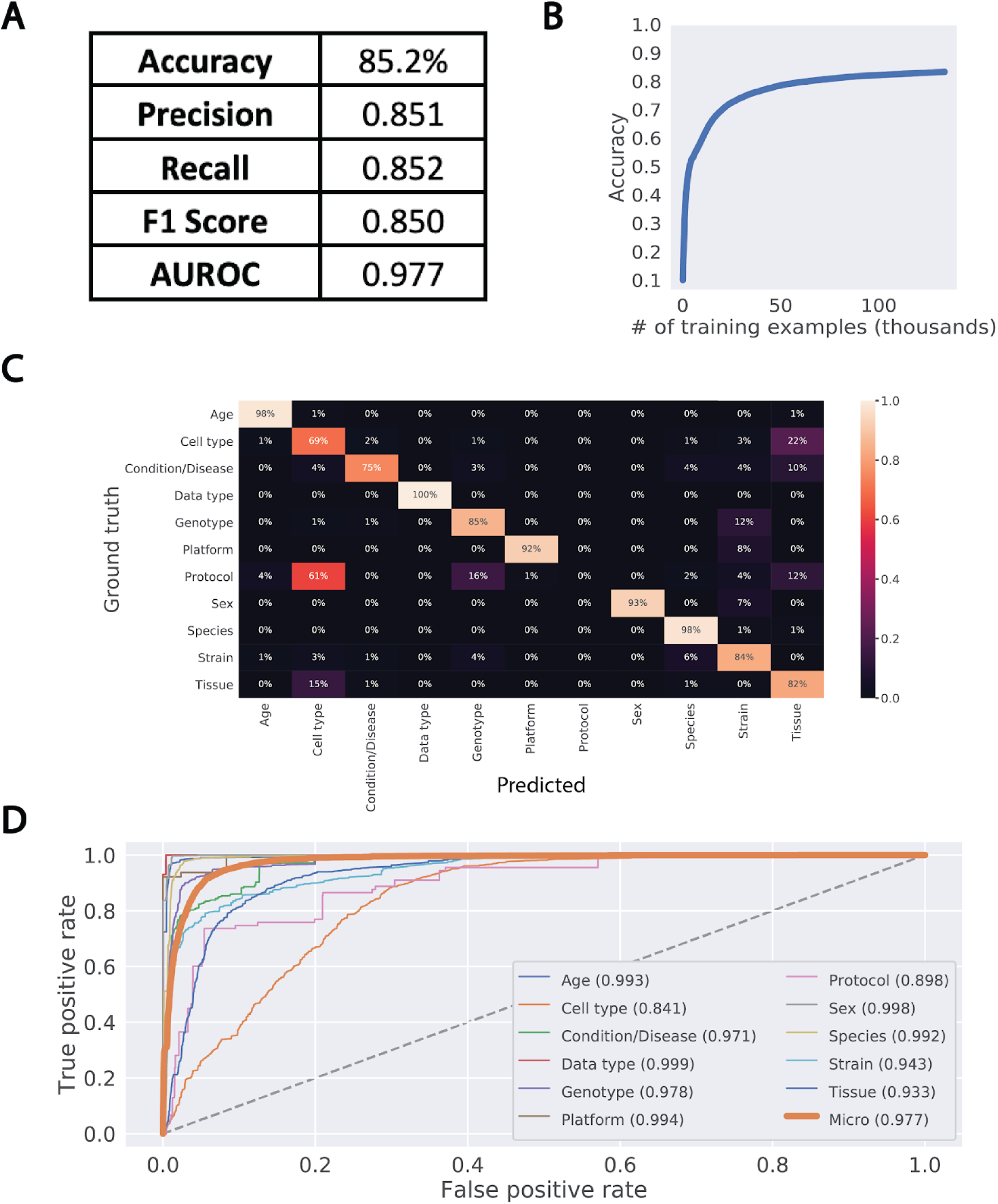
Performance of bi-LSTM in metadata category classification. **(A)** Accuracy, precision, recall, F1 score, and average AUROC calculated for all categories combined. **(B)** Accuracy of model classification on training set (y-axis) plotted against the number of training examples input (in thousands). **(C)** Percentage of each category correctly classified, shown as a heatmap, with predicted values on the x-axis and ground truth labels on the y-axis. **(D)** Receiver operating characteristic (ROC) curves for each category along with the average over all test set examples (micro average).

### High accuracy of metadata prediction observed for Species, Disease/Condition and Strain

We next applied this trained bi-LSTM to extract metadata from the TITLEs of samples withheld from classifier training and testing. For each metadata category, we selected up to 1,000 samples with TITLEs of at least 5 words that had a ground truth annotation for that category already present in BioSample. Using an n-gram based approach, NER was performed on these TITLEs, picking out the 11 metadata categories from n-grams of length 2 to 7 from this longer free text. Evaluation of the prediction algorithm was done by comparing any predicted metadata for each category to the ground truth annotations. Specifically, if the entirety of the predicted or ground truth metadata was a substring of the other, or if the cosine similarity between the two was greater than 0.7 in the embedding space, the prediction was considered correct. Evaluation of performance in this manner showed high accuracy for Condition/Disease, Data type, Sex, Species and Strain (Table 1). Moderate performance was noted for Genotype and Tissue, with lower than 50% accuracy seen in Platform, Age, and Cell type prediction.

Manual examination of predictions illustrated several patterns of incorrect predictions under our current evaluation criteria (Supplementary Table 4). Many incorrect predictions occurred when the model extracted metadata that didn’t match the concept of the category predicted. These *bona fide* incorrect predictions occurred most commonly on more heterogeneous fields like Cell type, Strain and Genotype. However, incorrect predictions from TITLEs that did not actually contain the exact ground truth value as a substring, but either contained a variation of the actual value or a completely different value that falls under the same metadata category, were also prevalent. In these cases, the model correctly selects a metadata category from the TITLE, but that prediction fails to match the underlying ground truth annotation. In many of these cases, an argument could be made that the model’s predicted annotation from the TITLE is a better descriptor of the metadata category for that sample than the ground truth (Supplementary Table 4). Furthermore, since the model predicts only 2-grams through 7-grams, ground truth annotations that were a single word were challenging and often incorrect. These values are especially prevalent in the Age category that the model showed very low accuracy on and in the Sex category that had only 4 predictions out of 190 TITLEs. Despite these issues, these results indicate that our model can accurately extract metadata annotations across multiple entity types.

## Discussion

In this work, we illustrate several issues with current annotations of SRA samples in BioSample and present a fully automated framework for increasing the coverage of some of the key metadata terms that describe these samples. We trained a deep neural network that was able to classify 11 metadata categories and achieved high classification performance on most categories. Prediction of key metadata fields from sample TITLEs using this model yielded high accuracy NER extractions of several categories, and also highlighted discrepancies between TITLEs of samples and underlying metadata.

User-defined fields, though useful and even necessary in certain situations, have led to a significant increase in heterogeneity across this dataset and others (5). The use of word embeddings for clustering attributes by semantic similarity revealed a lack of normalization in attribute naming, mostly in the form of small deviations in spelling and punctuation (e.g., cell type and Cell type). This clustering, along with the evaluation of predicted metadata, also highlighted that values under the same attribute often vary substantially in concept. The overall heterogeneity seems to be caused by a combination of user-input error and a lack of understanding of what values go under what attribute, even in BioSample defined attributes. The latter seems to be especially prevalent in defining a sample’s Cell type, Tissue, Strain and Genotype, as semantic analysis of those attributes showed they share a high degree of similarity in the embedding space. As a consequence of these issues, searches for specific metadata under a given attribute may lead to incorrect or incomplete returns for researchers, making it difficult to capitalize on the full potential of large resource collections such as the SRA.

Previous work has highlighted the utility of word embeddings in clustering metadata categories that share a high degree of similarity in the embedding space (18). We have shown that this clustering can also be utilized to improve the uniformity of SRA BioSample metadata and can be applied to increase class coverage for training an NER model. However, high levels of similarity between attributes can blur the distinction between some of the merged metadata categories and represents an important source of error in classification and subsequent prediction. Tokens outside of the embedding vocabulary (i.e., out of vocabulary words or OOVs), currently ignored by our model, also likely contributed to missed classifications and incorrect predictions. Using a similar framework with learned embeddings or with fine-tuned, pre-trained models like BERT (16) may better delineate the classification categories and limit OOV words. Furthermore, incorporating preprocessing steps such as part of speech tagging and stemming may limit noise and improve our prediction accuracy.

Further analysis of the incorrect and missing predictions may also bring forth algorithmic solutions for increasing prediction quality. Currently, the prediction algorithm only considers 2-grams through 7-grams, contributing to the low prediction numbers and accuracies for categories that contain a significant proportion of 1-grams (e.g., Sex and Age). However, consideration of 1-grams currently leads to many spurious metadata predictions, and more work is needed to determine how to better capture these classes with our model. Moreover, the variability in underlying ground truth metadata between and within categories creates a challenge in selecting an appropriate and rigorous evaluation approach. A more systematic and improved method for evaluation would likely elucidate algorithmic adjustments that would improve subsequent metadata prediction quality.

The publishing of FAIR principles in 2016 outlined a guide for proper data management practice and highlighted the need for findable, accessible, interoperable and reusable metadata in science (19). The large degree of heterogeneity inherent in biological data repositories makes the problems underlying the improvement of biomedical metadata quality non-trivial and illustrates that a one-size-fits-all solution is likely idealistic. The work presented here applies mainly to the reusability aspect of FAIR, focusing specifically on metadata plurality. Though our metadata prediction accuracy remains variable, we have shown that an NLP based methodology has the potential to augment current efforts to improve metadata completeness and quality. The automation of our pipeline provides a significant scaling advantage over manual curation and can easily be adapted for repositories that use similar attribute-value pair relationships in their metadata. It may be the case that this heterogeneity is a problem only completely solved by more careful annotation upon submission, but in its current form, our model represents a step towards improving SRA metadata plurality and reusability in the present.

## Contributions

Original concept: B.T., S.M., C.H.; experimental design: B.T., A.K., S.M., C.H., H.C.; data analysis: B.T., A.K. S.M.; implementation: B.T., A.K.; validation design: B.T., S.M., A.K.; data interpretation: B.T., A.K., S.M., D.S., M.D., C.H., H.C.; manuscript writing: A.K., B.T., H.C.

