## Supplementary Figures and Tables for "Increasing metadata coverage of SRA BioSample entries using deep learning based Named Entity Recognition"

1. **Supplementary Figure 1 -** Breakdown of training and testing data.
2. **Supplementary Figure 2** - *Homo sapien* and TITLE sample metadata coverage.
3. **Supplementary Figure 3** - Training set metadata categories show a large degree of similarity in the embedding space.
4. **Supplementary Table 1** - Selected attributes from BioSample and their corresponding category name for model training.
5. **Supplementary Table 2** - Cosine similarity between attributes used in merging.
6. **Supplementary Table 3** - Coverage increase for merged metadata categories.
7. **Supplementary Table 4** - Incorrect metadata predictions across all 11 categories.

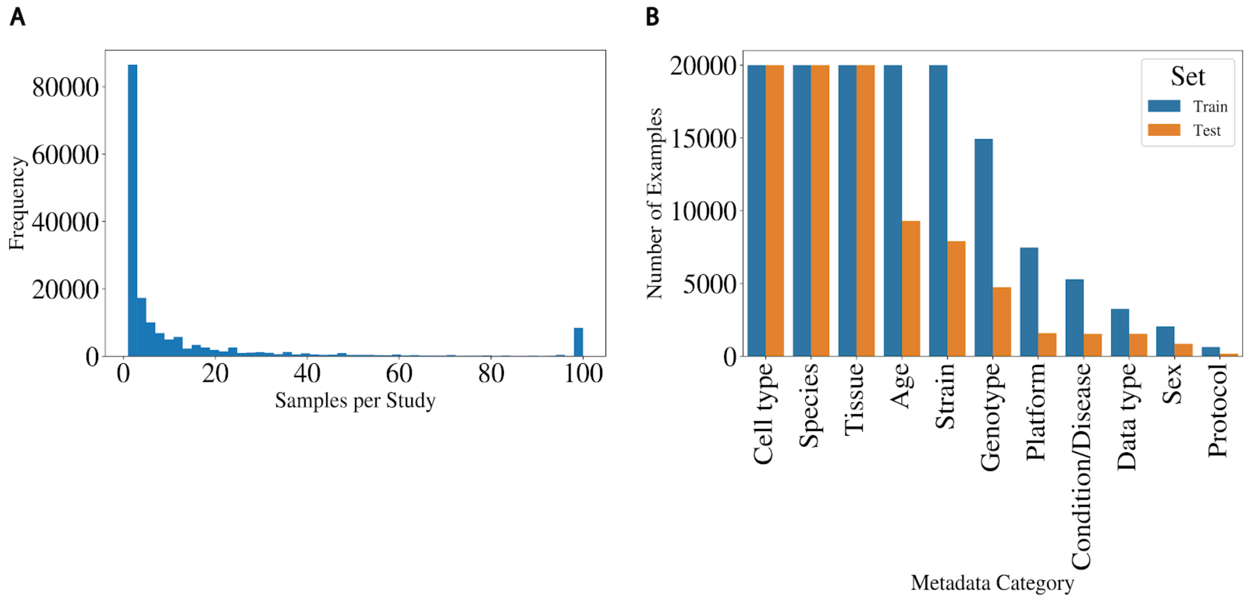


**Supplementary Figure 1.** Breakdown of training and testing data. **(A)** Histogram showing the distribution of samples in each study (capped at 100) used to generate training and test sets. **(B)** Number of training and test examples for each metadata category (capped at 20,000).

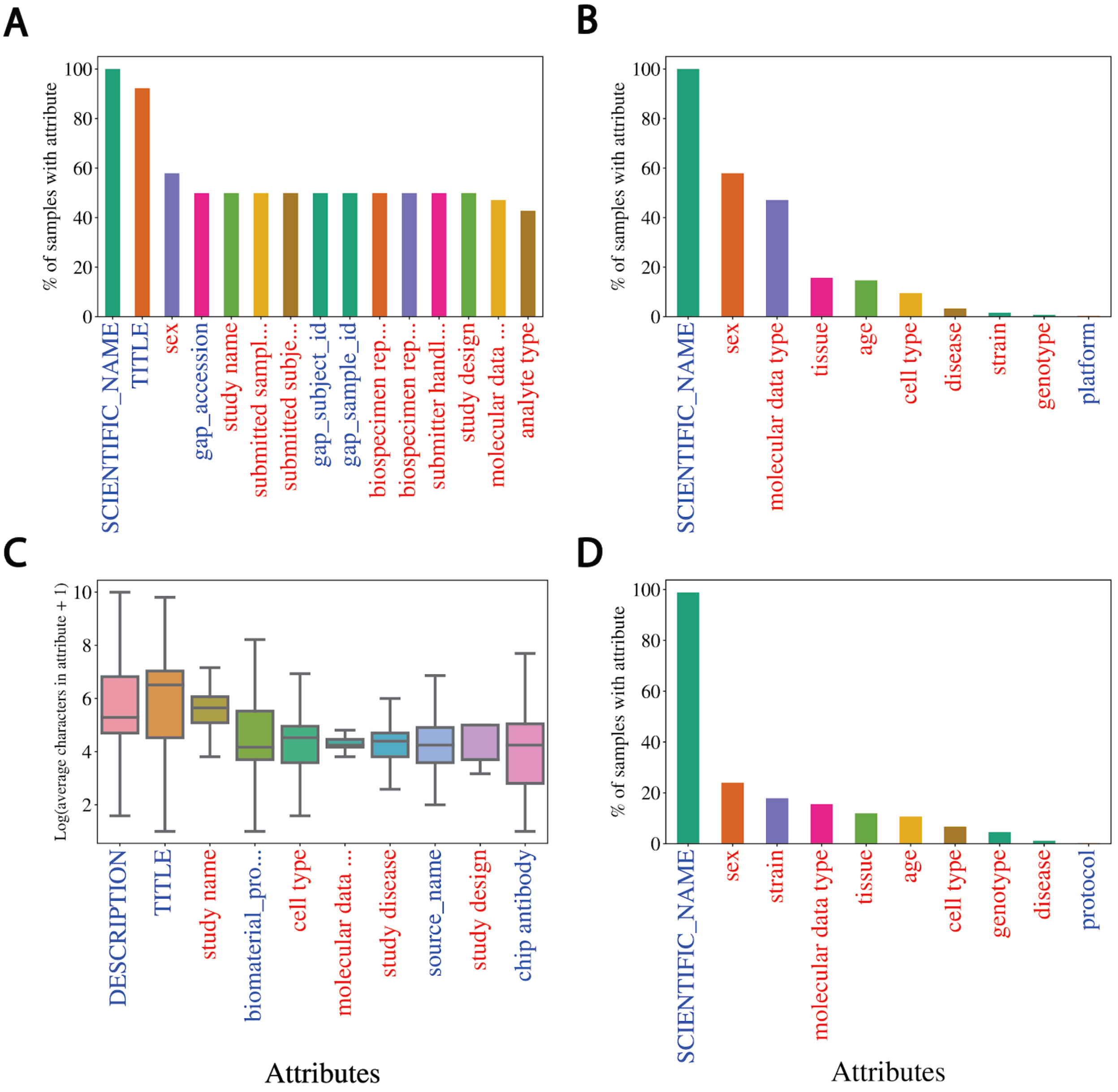


**Supplementary Figure 2.** *Homo sapien* and TITLE sample metadata coverage. **(A)** Top 15 attribute coverage, **(B)** 11 selected attribute coverage, and **(C)** top 10 distributions (by mean) of $Log_2$(average characters) for *Homo sapien* samples only. Samples annotated with "Homo sapien" show relatively low coverage for most attributes, but contain longer TITLEs for NER. **(D)** Selected attribute coverage for samples with a TITLE attribute only. This shows that samples with TITLEs are missing metadata in all attributes besides SCIENTIFIC_NAME.

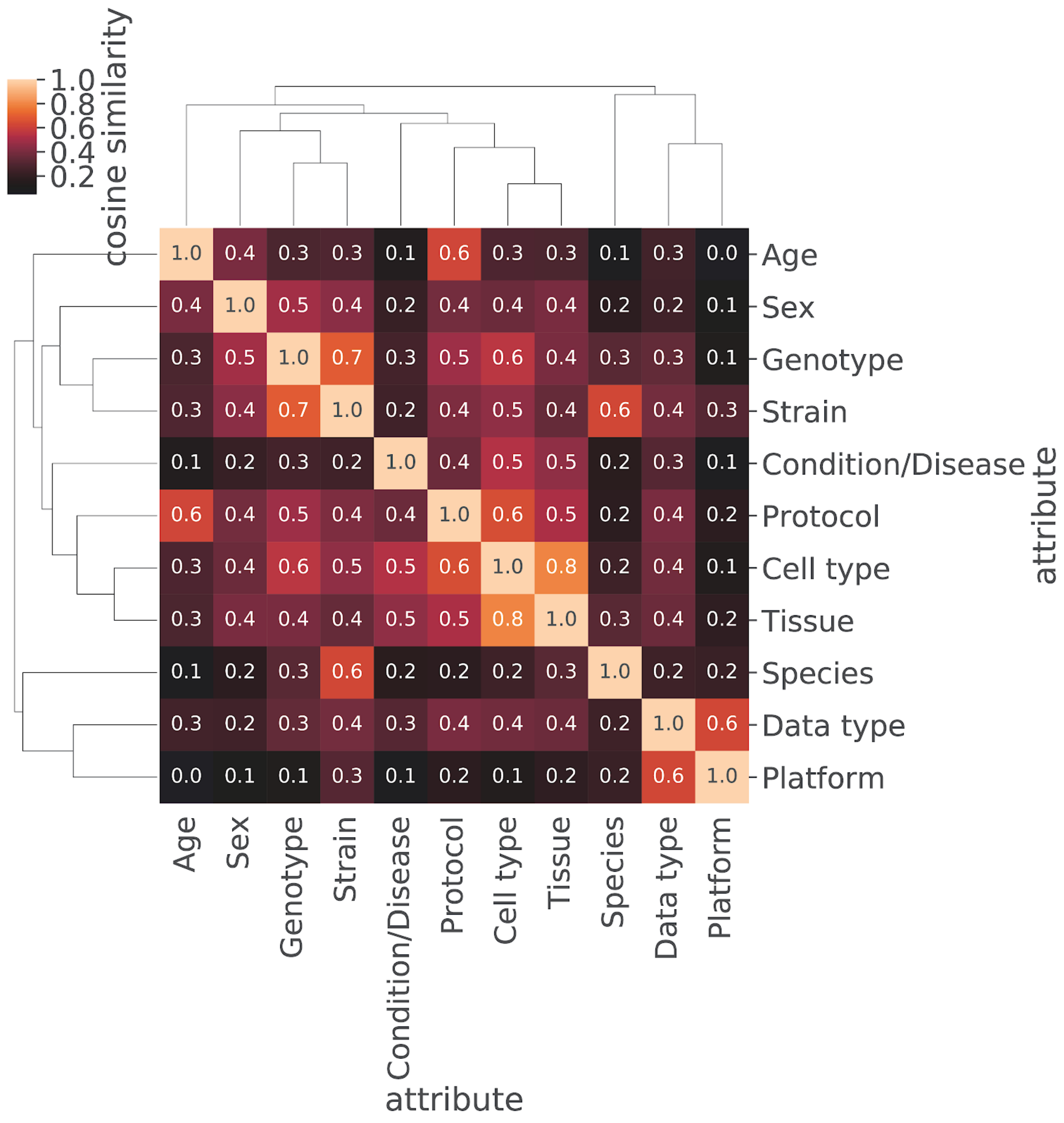


**Supplementary Figure 3.** Training set metadata categories show a large degree of similarity in the embedding space. Cosine similarity heatmap of mean vectors for the 11 categories in the training set with hierarchical clustering shown. Many categories in the training data show a high degree of semantic similarity in the embedding space.

**Supplementary Table 1.** Selected attributes from BioSample and their corresponding category name for model training. The category names represent multiple attributes that were merged with the selected attribute based on cosine similarity in the word embedding space.

| Selected Attribute | Category Name |
|---|---|
| age | Age |
| cell type | Cell Type |
| disease | Condition/Disease |
| molecular data type | Data type |
| genotype | Genotype |
| platform | Platform |
| protocol | Protocol |
| sex | Sex |
| SCIENTIFIC_NAME | Species |
| strain | Strain |
| tissue | Tissue |

**Supplementary Table 2**. Cosine similarity between attributes used in merging. Cosine similarities of mean embedding vectors for each attribute compared to selected attributes. The selected attributes used are bolded. All those attributes with a cosine similarity to the bolded attribute of greater than 0.8 were selected for merging and are shown here.

| Category Name | Attribute | Cosine Similarity |
|---|---|---|
| Species | **SCIENTIFIC_NAME** | 1.000 |
| | Organism | 0.917 |
| | host scientific name | 0.911 |
| | organism | 0.903 |
| | host_scientific_name | 0.886 |
| | host | 0.884 |
| | nat-host | 0.859 |
| | specific host | 0.853 |
| | host organism | 0.851 |
| | host species | 0.831 |
| | specific_host | 0.805 |
| | HostSpecies | 0.801 |
| Strain | **strain** | 1.000 |
| | strain background | 0.932 |
| | host_genotype | 0.931 |
| | background strain | 0.927 |
| | strain/background | 0.925 |
| | genetic background | 0.912 |
| | mouse strain | 0.906 |
| | Mouse_Strain | 0.899 |
| | StrainOrLine | 0.894 |
| | host strain | 0.891 |
| | stain | 0.891 |
| | host_strain | 0.881 |
| | strain or line | 0.880 |
| | strain name | 0.879 |
| | host infra-specific name | 0.870 |
| | host genotype | 0.865 |
| | background | 0.864 |
| | host_breed | 0.849 |
| | Strain | 0.836 |

| | | |
|---|---|---|
| | maternal strain | 0.835 |
| | paternal strain | 0.812 |
| Cell type | **cell type** | 1.000 |
| | cell_type | 0.897 |
| | source_name | 0.886 |
| | source cell type | 0.885 |
| | CellType | 0.869 |
| | progenitor cell type | 0.866 |
| | cell types | 0.865 |
| | cell-type | 0.853 |
| | cell description | 0.849 |
| | CELL_TYPE | 0.845 |
| | biomaterial_type | 0.841 |
| | tissue/cell type | 0.827 |
| Genotype | **genotype** | 1.000 |
| | genotype/variation | 0.940 |
| | Genotype | 0.895 |
| | mutant | 0.882 |
| | mutation | 0.847 |
| | plant genotype | 0.824 |
| | genetic variation | 0.812 |
| | idh2.gene.mutation | 0.801 |
| | host genotype | 0.801 |
| Condition/Disease | **disease** | 1.000 |
| | disease status | 0.833 |
| | tumor type | 0.828 |
| | health state | 0.828 |
| | cancer type | 0.826 |
| | cell description | 0.813 |
| Tissue | **tissue** | 1.000 |
| | tissue_type | 0.913 |
| | organism part | 0.848 |
| | tissue-type | 0.827 |
| | source_name | 0.800 |
| Sex | **sex** | 1.000 |
| | host_sex | 0.997 |
| | Sex | 0.996 |
| | host ex | 0.995 |
| | sex_infant_1 | 0.995 |

| | | |
|---|---|---|
| | sex / reassigned sex | 0.995 |
| | Host_Gender | 0.995 |
| | babygender_m_f | 0.994 |
| | gender | 0.994 |
| | ; | 0.987 |
| | breeding direction | 0.981 |
| | Gender | 0.939 |
| | host sex | 0.930 |
| | SEX | 0.900 |
| | sex_def_prob | 0.865 |
| Age | **age** | 1.000 |
| | Age | 0.832 |
| Data type | **molecular data type** | 1.000 |
| Platform | **platform** | 1.000 |
| | Platform | 0.942 |
| | instrument_model | 0.935 |
| | Sequencing_method | 0.934 |
| | sequencing method | 0.932 |
| | INSTRUMENT_MODEL | 0.930 |
| | SequencingTechnology | 0.926 |
| | sequencer | 0.916 |
| | illumina_technology | 0.914 |
| | labversion description | 0.914 |
| | illumina_technlogy | 0.910 |
| | Sequencer | 0.910 |
| | seq_meth | 0.903 |
| | sequencing_platform | 0.892 |
| | seq_methods | 0.891 |
| | sequencing_machine | 0.890 |
| | sequencing_method | 0.874 |
| | runchemistry | 0.850 |
| | seq_method | 0.802 |
| Protocol | **protocol** | 1.000 |
| | technology | 0.887 |
| | extract_protocol | 0.860 |
| | experiment type | 0.858 |
| | assay | 0.850 |
| | protocol description | 0.844 |
| | application | 0.843 |

| | | |
|---|---|---|
| | library_type | 0.800 |

**Supplementary Table 3.** Coverage increase for merged metadata categories

| Category Name | Selected Attribute | Selected Attribute Count | Category Name Count | Fold Increase |
|---|---|---|---|---|
| Age | age | 363161 | 370206 | 1.02 |
| Cell type | cell type | 150652 | 685002 | 4.55 |
| Condition/Disease | disease | 36349 | 40315 | 1.11 |
| Data type | molecular data type | 544515 | 544515 | 1.00 |
| Genotype | genotype | 117332 | 154376 | 1.32 |
| Platform | platform | 3476 | 147827 | 42.53 |
| Protocol | protocol | 5851 | 9290 | 1.59 |
| Sex | sex | 643290 | 808204 | 1.26 |
| Species | SCIENTIFIC_NAME | 2773124 | 3635784 | 1.31 |
| Strain | strain | 598496 | 730595 | 1.22 |
| Tissue | tissue | 446429 | 499537 | 1.12 |

**Supplementary Table 4.** Incorrect metadata predictions across all 11 categories

| Category Name | TITLE Sentence | Predicted | Actual |
|---|---|---|---|
| Species | 32% Streptococcus mutans UA159 69 | Streptococcus mutans | mixed sample |
| Species | 68% Lactobacillus casei 4646 | Lactobacillus casei | mixed sample |
| Species | A cucumber genomic variation map reveals impact human selection | A cucumber | cucumis sativus |
| Species | Bulk soils compared rhizosphere communities plant Saxifraga oppositifolia along glacier chronosequence | Saxifraga oppositifolia | unidentified bacterium |
| Species | CBo1 isolated gut Bactrocera oleae | Bactrocera oleae | chryseobacterium sp. 113(2015) |
| Species | Endophytic isolate branches Citrus sinensis | Citrus sinensis | curtobacterium sp. er1/6 |
| Species | Environmental/Metagenome sample subarctic root fungi | subarctic root | fungi |
| Species | Generic sample Fusobacterium nucleatum | Fusobacterium nucleatum | fusobacterium phage funu1 |
| Species | Generic sample Homo sapiens respiratory syncytial virus | Homo sapiens | human orthopneumovirus |
| Species | Generic sample Mimulus guttatus | Mimulus guttatus | erythranthe guttata |
| Species | Human respiratory syncytial virus | syncytial virus | human orthopneumovirus |
| Species | Hyphomicrobium facile subsp | facile subsp | hyphomicrobium facile |
| Species | Ip394 Japanese crested ibis (founder E) | Ip394_Japanese crested ibis | nipponia nippon |
| Species | Lathyrus pratensis clone 275 reared Lathyrus pratensis | Lathyrus pratensis | acyrthosiphon pisum |
| Species | MIMARKS Survey related sample Borrelia burgdorferi species complex | Borrelia burgdorferi | borrelia sp. s07cg098 |
| Species | MIMARKS Survey related sample Borrelia burgdorferi species complex | Borrelia burgdorferi | borrelia sp. s07as020 |
| Species | MIMARKS Survey related sample Borrelia burgdorferi species complex | Borrelia burgdorferi | borrelia sp. s10cg023 |
| Species | MIMARKS Survey related sample Borrelia burgdorferi species complex | Borrelia burgdorferi | borrelia sp. s08cg103 |
| Species | Medicago truncatula | Medicago truncatula | mixed culture |
| Species | Mycobacterium lepromatosis red squirrel M1960/270411 | squirrel M1960/270411 | mycobacterium lepromatosis |
| Species | Phylogenomic characterization causative strain one largest worldwide outbreaks Legionnaires' disease occurred Portugal 2014 | characterization causative | legionella pneumophila |
| Species | Picea engelmannii cultivated The Arboretum Horsholm University Copenhagen Denmark | Picea engelmannii | populus tremula |
| Species | Plant DNA metabarcoding data uncleaned surface cleaned wood subfossil tree trunks | subfossil tree | pinus sylvestris |
| Species | Plant DNA metabarcoding data uncleaned surface cleaned wood subfossil tree trunks | subfossil tree | pinus sylvestris |
| Species | Plant DNA metabarcoding data uncleaned surface cleaned wood subfossil tree trunks | subfossil tree | pinus sylvestris |
| Species | Root tissue samples Caladium hortulanum (MissMuffet) | Caladium hortulanum | caladium bicolor |
| Species | SYNE2/ESR2A CALM1 loci Clupea pallisii PH3 | Clupea pallisii | clupea pallasii |
| Species | SYNE2/ESR2A CALM1 loci Clupea pallisii PH5 | Clupea pallisii | clupea pallasii |
| Species | Small RNAs Miscanthus rhizomes | Miscanthus rhizomes | miscanthus x giganteus |
| Species | Stable isotope probing targeting methane-utilizing bacteria arctic lake sediment | arctic lake | bacteria |
| Species | Streptococcus agalactiae | Streptococcus agalactiae | mixed culture |

| | | | |
|---|---|---|---|
| Species | Transcriptomics potato tubers infected Streptomyces turgidiscabies | Streptomyces turgidiscabies | solanum tuberosum |
| Species | Uncultivated Candidatus UAP2 archaeon UBA543 genome recovered ERX552263 | Candidatus UAP2 | archaeon uba543 |
| Species | Whole genome analysis Clostridium saccharogumia VE202-01 | Clostridium saccharogumia | [clostridium] saccharogumia |
| Species | cecal microbiota sample Microtus brandti | Microtus brandti | lasiopodomys brandtii |
| Species | drifting hotspot microbial diversity | microbial diversity | caulerpa taxifolia |
| Species | foliar sample Castanea sativa | Castanea sativa | uncultured fungus |
| Species | foliar sample Vitis vinifera | Vitis vinifera | uncultured fungus |
| Species | foliar sample Vitis vinifera | Vitis vinifera | uncultured fungus |
| Species | hRSV RSV+PIV3 F insert C 03 | hRSV RSV+PIV3 | human orthopneumovirus |
| Species | live-1 pig low backfat thickness | live-1 pig | sus scrofa |
| Species | sativus contaminated Alternaria brassicicola Abra43 | Alternaria brassicicola | raphanus sativus var. sativus |
| Species | treated plants polyethylene glycol PEG NaCl carried transcriptomic metabolomics measurements across time-course five days | polyethylene glycol | calotropis procera |
| Species | unspecified weed species yellowing mosaic-like symptoms | weed species | unidentified plant |
| Strain | 1 New Tech Library sample 21095 | New Tech | 2.4.1 |
| Strain | 4 mM - R-1 (ncra) | R-1 (ncra) | fgsc 2489 |
| Strain | A-1 rbs | A-1 rbs | tc1769 |
| Strain | Aspergillus niger NRRL 3 delta araR Preculture Gene expression profiling - Pa | NRRL 3 | nrrl3 delta arar |
| Strain | Aureus strain DSM799 | strain DSM799 | dsm 799 |
| Strain | CR0044 serovar Typhi 13438 | CR0044 serovar | 13438 |
| Strain | CRISPR 128 genomic zebrafish loci ‚Äì sgRNAs ‚Äì Pull-down expt ‚Äì Canonical ‚Äì Mutant pool1 | expt ‚Äì | tu |
| Strain | Candida glabrata genome-wide nucleosome data Jan2809 | data Jan2809 | clib 138 |
| Strain | CgPho2 ChIP Pi media w/ CgPho4 | w/ CgPho4 | cg40 |
| Strain | Escherichia coli serovar O101:H33 175562 | serovar O101:H33 | 175562 |
| Strain | Escherichia coli serovar O11:H25 204427 | serovar O11:H25 | 204427 |
| Strain | Fungi PEN-treated leaves SM2 rep | SM2 rep | 49 |
| Strain | Generic sample Enterococcus columbae DSM 7374 = ATCC 51263 | ATCC 51263 | dsm 7374 |
| Strain | Khuskia oryzae ATCC28132 Annotated Standard Draft | ATCC28132 Annotated | atcc 28132 |
| Strain | LES CF sputum CF03 isolate 14 | CF03 isolate | liverpool epidemic strain |
| Strain | LPS 48h pf 13 Replicate 1 | 13 Replicate 1 | c57bl/6babr |
| Strain | Louis clinical isolate (#12) | Louis clinical | 68 |
| Strain | Neurospora crassa FGSC2489 X | FGSC2489 X | 2489x8790 370a |
| Strain | Neurospora crassa FGSC2489 X | FGSC2489 X | 2489x8790 90a |
| Strain | Neurospora crassa FGSC2489 X | FGSC2489 X | 2489x8790 305a |
| Strain | Neurospora crassa FGSC2489 X | FGSC2489 X | 2489x8790 362a |
| Strain | Pyronema omphalodes CBS 100304 | CBS 100304 | cbs100304 |
| Strain | RNA-seq data Paratyphi A 45157 grown stationary phase 42C | A 45157 | paratyphi a |
| Strain | Resequencing evolved clone P4C7 | clone P4C7 | cc2937 |
| Strain | Salmonella enterica enterica serovar Salmonella Monschaui 369918 | Salmonella Monschaui | 369918 |
| Strain | Salmonella enterica serovar Agona SL483 uid59431 serovar Agona 9165 | SL483 uid59431 | 9165 |
| Strain | Salmonella enterica subsp I enterica serovar Ajiobo 63502 | serovar Ajiobo | 63502 |
| Strain | Salmonella enterica subsp I enterica serovar Goelzau H120240584 | serovar Goelzau | h120240584 |
| Strain | Salmonella enterica subsp I enterica serovar | serovar Monschaui | 63893 |

| | | | |
|---|---|---|---|
| | Monschaui 63893 | | |
| Strain | Salmonella enterica subsp I enterica serovar Stanley 32479 | serovar Stanley | 32479 |
| Strain | Salmonella enterica subsp I enterica serovar Stanley 78659 | serovar Stanley | 78659 |
| Strain | Sample UHHS A | UHHS A | abuh434 |
| Strain | Use samples cell lines derived common laboratory model organisms | laboratory model | lvpib12 |
| Strain | VC1639 mutant polymyxin B replicate 2 | polymyxin B | vc1639 |
| Strain | Wildtype head X1 single cell 33 | X1 single | asexual ciw4 |
| Strain | aeruginosa PASGNDM strains isolated Singapore | PASGNDM strains | pasgndm593 |
| Strain | hot springs Yunnan Tibet | Yunnan Tibet | a12b |
| Strain | military healthcare system | military healthcare | mrsn7485 |
| Strain | military healthcare system | military healthcare | mrsn7087 |
| Strain | military healthcare system | military healthcare | mrsn15047 |
| Strain | military healthcare system | military healthcare | mrsn7132 |
| Strain | military healthcare system | military healthcare | mrsn7376 |
| Strain | military healthcare system | military healthcare | mrsn11693 |
| Strain | military healthcare system | military healthcare | mrsn11754 |
| Strain | military healthcare system | military healthcare | mrsn7237 |
| Strain | military healthcare system | military healthcare | mrsn7498 |
| Strain | military healthcare system | military healthcare | mrsn908 |
| Strain | military healthcare system | military healthcare | mrsn7730 |
| Strain | military healthcare system | military healthcare | mrsn953 |
| Strain | military healthcare system | military healthcare | mrsn7188 |
| Strain | military healthcare system | military healthcare | mrsn3384 |
| Strain | military healthcare system | military healthcare | mrsn7140 |
| Strain | viewpoint 155 | viewpoint 155 | c57bl/6j & 129s5svevbrd |
| Cell type | 14SA cells | 14SA cells | fetal muscle myoblast |
| Cell type | 3-HA ChIP-seq ES cells 4dRA 1hr DOX | 4dRA 1hr | mesc 4dra |
| Cell type | 34516 mESC 1h chase Rep 1 | 34516 mESC | mouse embryonic stem (mes) cells |
| Cell type | 34521 mESC 3h chase Rep 3 | 34521 mESC | mouse embryonic stem (mes) cells |
| Cell type | 37054 mESC wt 12h pulse | 37054 mESC | mouse embryonic stem (mes) cells |
| Cell type | 37061 mESC Mettl3KO #2 3h pulse | mESC Mettl3KO | mouse embryonic stem (mes) cells |
| Cell type | AR ChIP sequencing 2hr 100nM DHT treated VCaP cells | VCaP cells | prostate cancer cells |
| Cell type | Activated Erk1/2 Promotes Formation Chromatin Features Inherent Developmental Promoters Mouse Embryonic Stem Cells | Stem Cells | es cells |
| Cell type | Aorta smooth muscle cells AoSMC2 | cells AoSMC2 | aorta smooth muscle cells |
| Cell type | ChIP-Seq Med1 mES cells | mES cells | v6.5 embryonic stem cells |
| Cell type | ChIP-Seq Smc3 mES cells | mES cells | v6.5 embryonic stem cells |
| Cell type | ChIP-Seq analysis H3K79me2 mouse ESCs shControl knockdown treatment replicate 2 | analysis H3K79me2 | mouse embryonic stem cells |
| Cell type | ChIP-Seq analysis HA-Phf5a mouse ESCs doxycycline treatment replicate 1 | mouse ESCs doxycycline treatment | mouse embryonic stem cells |
| Cell type | ChIP-Seq analysis Leo1 mouse ESCs shControl knockdown treatment replicate 1 | mouse ESCs shControl knockdown treatment | mouse embryonic stem cells |
| Cell type | Chromatin IP ERŒ± MCF-7 cells transfected LRH-1 siRNA Replicate 1 | ERŒ± MCF-7 | breast cancer |
| Cell type | Chromatin IP HA MCF-7 cells transfected HA-LRH1 vector Replicate 2 | MCF-7 cells | breast cancer |
| Cell type | DDX5-/- mESCs Line #1 (miRNA-seq) | DDX5-/- mESCs | mus musculus |
| Cell type | DDX5-/- mESCs Line #2 (miRNA-seq) | DDX5-/- mESCs | mus musculus |

| | | | |
|---|---|---|---|
| Cell type | DKO1 H3K27ac replicate 1 ChIP-seq | DKO1 H3K27ac | human colorectal cancer cell line |
| Cell type | DS20201 SE DNaseI-seq seed coat cell 4DPA | cell 4DPA | seed coat cell (4 days past anthesis) |
| Cell type | ENCODE biosample ENCBS478HYO: Differentiated 3T3-L1 (Day 8) | Differentiated 3T3-L1 | adipocyte |
| Cell type | ENCODE biosample ENCBS513ENC: neurons derived H1 embryonic stem cells | stem cells | neural cell |
| Cell type | Embryonic Stem cells Jarid2 -/- rescue Jarid2K116A BAC H3K27me3 IP | Stem cells | esc serum + lif grown |
| Cell type | Enteroendocrine Cells RNA-seq biological rep1 | Enteroendocrine Cells | adult epitheial cells; proximal 1/3 of intestine |
| Cell type | GRO-Seq analysis mouse ESCs flavopiridol treatment replicate 2 | analysis mouse ESCs flavopiridol treatment | mouse embryonic stem cells |
| Cell type | Gm6871-HA ChIP-seq mES cells | mES cells | embryonic stem cells |
| Cell type | H3K27ac marks na√Øve LIS2 ES stem cells | stem cells | escs |
| Cell type | H3K27ac marks primed LIS2 ES stem cells | stem cells | escs |
| Cell type | H3K27ac marks primed WIS2 ES stem cells | stem cells | escs |
| Cell type | H3K27me3 ChIP | H3K27me3 ChIP | fibroblasts |
| Cell type | H3K27me3 marks primed WIS2 ES stem cells | stem cells | escs |
| Cell type | H3K4me1 sequencing non-treated (DMSO vehicle) MCF-7 cells | MCF-7 cells | breast cancer cells |
| Cell type | H3K4me3 HFFs H3K27ac HFFs input | HFFs H3K27ac | foreskin fibroblast |
| Cell type | H3K4me3 marks na√Øve WIS2 ES stem cells | stem cells | escs |
| Cell type | H3K9me3 marks na√Øve C1 induced stem cells | stem cells | ips cells |
| Cell type | H3K9me3 marks na√Øve WIBR3 embryonic stem cells | stem cells | hes cells |
| Cell type | H3K9me3 marks primed WIBR3 embryonic stem cells | stem cells | hes cells |
| Cell type | H929 cells expressing FAM46CWTGFP 72 hrs rep3 total RNA | H929 cells | multiple myeloma cell line |
| Cell type | HSC cd41low cd9low Gfi1 KO | HSC cd41low cd9low | hematopoietic stem cells cd41low cd9low |
| Cell type | HT29 10 ŒºM 5-Aza | ŒºM 5-Aza | colon cancer |
| Cell type | HUVEC AX15839 treated cells replicate 2 | HUVEC AX15839 | huvec cells |
| Cell type | HUVEC AX15839 treated cells replicate 3 | HUVEC AX15839 | huvec cells |
| Cell type | HUVEC PPARŒ≤/Œ¥ DMSO 24hr normoxia replicate 1 | PPARŒ≤/Œ¥ DMSO | human umbilical vein cells |
| Cell type | Input PolII Activin 1h ChipSeq biological replicate 2 | PolII Activin | embryonic teratoma |
| Cell type | LNCaP GIPZ EtOH AR ChIP m77 | ChIP m77 | prostate tumor |
| Cell type | LNCaP shGATA2 R1881 (10nM | LNCaP shGATA2 | prostate tumor |
| Cell type | Luciferase knockdown PC3 cells using shRNa replicate 2 | PC3 cells | prostate cancer cells |
| Cell type | MC1-ZE7 cells (Em+ high fraction) Hpa II | MC1-ZE7 cells | facs sorted emerald+ es cells |
| Cell type | Mettl3-KO EBs 0h Actinomycin treatment | Mettl3-KO EBs | embroid bodies |
| Cell type | Mettl3-KO ESCs 0h Actinomycin treatment | Mettl3-KO ESCs | embryonic stem cells |
| Cell type | Mmu bRG H3K27me3 replicate 1 | bRG H3K27me3 | basal radial glia |
| Cell type | PR1 - Pol2 mES cells + DMSO 18 hr ChipSeq | mES cells | v6.5 embryonic stem cells |
| Cell type | RNA-Seq analysis CEM cells upon treatment short hairpin UTX replicate 2 | analysis CEM cells upon treatment | human t cell leukemia cells |
| Cell type | RNA-Seq analysis HCT116 FBXW7 KO cells recovery heat shock treatment replicate 1 | analysis HCT116 | human colon cancer cells |
| Cell type | RNA-seq sample timecourse siRNA knock-down Tcf7l2: siRNA - Scrambled | timecourse siRNA | hepatocytes |
| Cell type | S2 T1 H3K27ac ChIP rep1 | T1 H3K27ac | s2 cell |
| Cell type | Sample 9 P14 effector CTLs IL-2 #1 | CTLs IL-2 | in vitro generated ctls |

| | | | |
|---|---|---|---|
| Cell type | Strains used generate cells must carry Fv2 sensitive allele (Fv2s) | generate cells | leukemia stem cell |
| Cell type | SuRE K562 cDNA | SuRE K562 | erythroleukemia |
| Cell type | Total RNA Pol2 ChIP-seq mES cells shSpt4 flavopiridol | cells shSpt4 | v6.5 embryonic stem cells |
| Cell type | Treg CNS2-/- Foxp3 low rep1 | CNS2-/- Foxp3 | regulatory t (treg) cells |
| Cell type | bisfulfite treated genomic DNA G1 fibroblasts A | DNA G1 | primary dermal fibroblasts |
| Cell type | bisfulfite treated genomic DNA arrested | DNA arrested | primary dermal fibroblasts |
| Cell type | human KRAS Q61H mutant form overexpressed HMEC cells [KRAS_Q61H-5] | HMEC cells | human mammary epithelial (hmec) cells |
| Cell type | long term repopulating hematopoietic stem cells control 3 | stem cells | bone marrow cells |
| Cell type | mESC m6A IP sample | mESC m6A | mouse embryonic stem cells |
| Cell type | nucARRB1 C4-2(high nuclear ARRB1)_H3K4me3 ChIP | ARRB1)_H3K4me3 ChIP | immortalised prostate epithelial cells |
| Cell type | overexpressed HMEC cells GFP18 control 10 | cells GFP18 control 10 | human mammary epithelial cells |
| Cell type | overexpressed HMEC cells GFP18 control 7 | cells GFP18 control 7 | human mammary epithelial cells |
| Cell type | overexpressed HMEC cells GFP30 control 4 | cells GFP30 control 4 | human mammary epithelial cells |
| Cell type | overexpressed HMEC cells GFP30 control 6 | cells GFP30 control 6 | human mammary epithelial cells |
| Cell type | overexpressed HMEC cells RAF1 overexpressed 5 | cells RAF1 overexpressed 5 | human mammary epithelial cells |
| Cell type | overexpressed HMEC cells 3 | HMEC cells 3 | mammary epithelial cells |
| Cell type | overexpressed HMEC cells 3 | HMEC cells 3 | mammary epithelial cells |
| Cell type | overexpressed HMEC cells 5 | HMEC cells 5 | mammary epithelial cells |
| Cell type | peripheral blood CD34+ cells pooled CD34pB | cells pooled CD34pB | peripheral blood cd34+ cells |
| Genotype | 1 cKO rep2 LPS + IL-4 | #NAME? | spi1fl/fl cd23t/+ |
| Genotype | 14028s ŒîSTM14 3463 + pBAD24 -2 | ŒîSTM14 3463 + | pbad24 |
| Genotype | 3 DMSO me3 rep2 | me3 rep2 | h3wt |
| Genotype | An WT No Carbon #1 | WT No | wildtype |
| Genotype | BLESS control sample mouse bone marrow | BLESS control | ssb1fl/flssb2fl/fl |
| Genotype | ChIP Kmg wild-type rep1 | Kmg wild-type | w[1118] |
| Genotype | Chlamydomonas reinhardtii 2137 - Copper sufficient - TAP - 4 | sufficient - | wt |
| Genotype | Chlamydomonas reinhardtii 2137 - Copper sufficient - TAP - 5 | sufficient - | wt |
| Genotype | D2 Control no2 | D2 Control | myoiats control |
| Genotype | Dietary emulsifiers directly alter human microbiota composition gene expression ex vivo potentiating intestinal inflammation | gene expression | c57bl6 |
| Genotype | Drosophila melanogaster - Ovary stages 1-8 follicle cell nuclei - OR s8 - Wild type - MNase-Seq | Wild type | modencode |
| Genotype | Ezh2 +/- replicate1 batch2 | Ezh2 +/- | sclcre |
| Genotype | Ezh2 -/- replicate6 | Ezh2 -/- | sclcre jak2v617f |
| Genotype | Gp 3 WT L 3 | WT L | sc5314 |
| Genotype | H4K8ac OR Head Nuclei D | H4K8ac OR | wild type |
| Genotype | HWT Start-seq females NextSeq rep2 | NextSeq rep2 | hwt |
| Genotype | HWT Start-seq mixed NextSeq rep2 | NextSeq rep2 | hwt |
| Genotype | Human set 4 36N H3K9Ac | 36N H3K9Ac | gfi1 36n/36s |
| Genotype | Input ChIP Aly-HA w1118 rep1 (negative control | (negative control | w[1118] |
| Genotype | Knockout Rv0954 H37Rv | Knockout Rv0954 | rv0954 ko |
| Genotype | M202 - control - rep 4 | control - rep | m202 |
| Genotype | Msx1-2 (f-f) WT - LCM RNA-Seq - Stroma | WT - | control (f/f) |
| Genotype | NPC Suv39h dn H3K9me3 ChIP-seq | Suv39h dn | suv39h double null |
| Genotype | P0testis mRNA seq: WT | P0testis mRNA seq: | wt, biological replicate 2, |

| | | | |
|---|---|---|---|
| | | WT | read1 |
| Genotype | P0testis mRNA seq: WT | P0testis_mRNA_seq: WT | wt, biological replicate 1, read1 |
| Genotype | PET111GFP EZ 6DO rep 1 | 6DO rep | pet111:gfp |
| Genotype | PET111GFP MZ 6DO rep 1 | 6DO rep | pet111:gfp |
| Genotype | PET111GFP MZ 7DO rep 1 | 7DO rep | pet111:gfp |
| Genotype | Productive VB18 allele PP PNA low B cells VB18 passenger mice | VB18 allele | vb18 passenger mice |
| Genotype | Productive VB18 allele Splenic GC B cells bglobin passenger mice | VB18 allele | bglobin mice |
| Genotype | RNA pol II ChIP-seq WT Rep2 | WT Rep2 | wildtype |
| Genotype | RNAseq round 2 atmorc4/7 rep 1 | atmorc4/7 rep | atmorc6 |
| Genotype | Ribosome protected fragments MZdicer 6hpf-2 | MZdicer 6hpf-2 | mzdicer mutant |
| Genotype | Whole testis RNA Dnmt3C IAP/WT 20 dpp 2 | Dnmt3C IAP/WT | heterozygous control |
| Genotype | Wild Type 6 month Female Heart mRNA rep1 | mRNA rep1 | wild type |
| Genotype | Wt HSC treated IFNŒ≥+VN-4 | treated IFNŒ≥+VN-4 | wild type |
| Genotype | anti-H3 T11ph Wild type culture #1 4h | T11ph Wild | wild type (mata/matalpha, ho::lys2/", lys2/", leu2::hisg/", ura3/") |
| Genotype | col0 - H3K9ac flg treated | H3K9ac flg | wt |
| Genotype | hebibaAOC Mock Time 24 hpt | hebibaAOC Mock | mutant |
| Genotype | htz1,àÜHYG Rad5+ t2 batch 3 | htz1,àÜHYG Rad5+ | htz1 delta |
| Genotype | iPSC 7dupASD1-C2 (GDB-Cf G) GTF2I KD | GTF2I KD | 7dupasd |
| Genotype | maize ear V12 rep 3 | V12 rep | b73 |
| Genotype | maize ear V14 drought rep 2 | drought rep | b73 |
| Genotype | maize leaf V14 drought rep 3 | drought rep | b73 |
| Genotype | maize leaf V14 drought rep 4 | drought rep | b73 |
| Genotype | maize tassel R1 rep 3 | R1 rep | b73 |
| Genotype | maize tassel V12 drought rep 2 | drought rep | b73 |
| Genotype | repeat 2 [Rlim KO] | [Rlim KO] | wt |
| Genotype | wild type retina sample 2 (WR2) | wild type | wildtype |
| Condition/Disease | inflamed region patient B1/non-stricturing | patient B1/non-stricturing | crohn's disease |
| Tissue | 4-6h afer egg laying | egg laying | embryonic |
| Tissue | 5 skin extract | skin extract | adult skin |
| Tissue | 6127 sigmoid stroma phase 1 | sigmoid stroma | colon |
| Tissue | Barin tissue sample - rep3 | tissue sample | brain |
| Tissue | CD Female Macroscopic inflammation Deep Ulcer (CCFA Risk 143) | Deep Ulcer | ileal biopsy |
| Tissue | CD Female Macroscopic inflammation No Deep Ulcer (CCFA Risk 018) | Deep Ulcer | ileal biopsy |
| Tissue | CD Male Macroscopic inflammation Deep Ulcer (CCFA Risk 150) | Deep Ulcer | ileal biopsy |
| Tissue | CD Male Macroscopic inflammation Deep Ulcer (CCFA Risk 216) | Deep Ulcer | ileal biopsy |
| Tissue | CD Male Macroscopic inflammation No Deep Ulcer (CCFA Risk 134) | Deep Ulcer | ileal biopsy |
| Tissue | CD Male Microscopic inflammation No Deep Ulcer (CCFA Risk 060) | Deep Ulcer | ileal biopsy |
| Tissue | Chandelier cells deep layer WT brain (CHC2-3) | deep layer | frontal cortex layers 5 & 6 |
| Tissue | DNA plasma patient P50_006 prostate cancer radical prostatectomy | P50 006 prostate | plasma before radical prostatectomy. |
| Tissue | DNA urine patient P50_008 prostate cancer radical prostatectomy | P50 008 prostate | urine after radical prostatectomy. |
| Tissue | DNA urine patient P50_010 prostate cancer radical prostatectomy | P50 010 prostate | urine before radical prostatectomy. |
| Tissue | Diet: High Fat | High Fat | pancreatic islets |
| Tissue | Diet: High Fat | High Fat | pancreatic islets |

| | | | |
|---|---|---|---|
| Tissue | Diet: High Fat | High Fat | pancreatic islets |
| Tissue | Diet: High Fat | High Fat | pancreatic islets |
| Tissue | Does timing matter pesticide resistance? One splice form variant MDR49 provides early | Does timing matter | whole body |
| Tissue | Drosophila simulans male body small RNA | body small | abdomen and thorax |
| Tissue | ENCODE biosample ENCBS046IMM: Heart embryonic 11 | biosample ENCBS046IMM: | heart |
| Tissue | ENCODE biosample ENCBS504VDB: Esophagus mucosa tissue aliquot received Gingeras lab conduct RNA-Seq assays | mucosa tissue | esophagus squamous epithelium |
| Tissue | ENCODE biosample ENCBS765HGS: EnTEX: Colon - Transverse | biosample ENCBS765HGS: | transverse colon |
| Tissue | ENCODE biosample ENCBS802FTI: Midbrain 88 embryonic 11 | biosample ENCBS802FTI: | midbrain |
| Tissue | ENCODE biosample ENCBS836VES: Skin lower leg aliquot received Gingeras lab conduct RNA-Seq assays | ENCBS836VES: Skin | lower leg skin |
| Tissue | ENCODE biosample ENCBS926XRV: Skin - Sun Exposed (Lower leg) | ENCBS926XRV: Skin | lower leg skin |
| Tissue | Enteropathy-associated T-cell Lymphoma tumor tissue | tumor tissue | eatl |
| Tissue | Fall'12 Day 42 Control Ross Heart Fat 1210 | Heart Fat | cardiac adipose |
| Tissue | Flag Leaf Tissue Biological Rep 2 | Leaf Tissue | flag leaf |
| Tissue | Generic sample Ctenopharynodon idellus | sample Ctenopharynodon | gill |
| Tissue | Generic sample Ctenopharynodon idellus | sample Ctenopharynodon | gill |
| Tissue | Generic sample Neohirasea fruhstorferi | sample Neohirasea | head and thorax |
| Tissue | Generic sample Oreophoetes peruana | sample Oreophoetes | head and thorax |
| Tissue | Genetic variations rectal cancer | variations rectal | primary tumor |
| Tissue | H3K27ac Chimpanzee Brain OccipitalPole PT1 | Brain OccipitalPole | occipital pole |
| Tissue | Hosui leaf defoliation research | leaf defoliation | flower buds |
| Tissue | Human sample Homo sapiens UUS2 tumor tissue | UUS2 tumor | uterus |
| Tissue | Invertebrate sample Acropora humilis: Tank - biological replicate 1 -noon- new moon | new moon | branch tip |
| Tissue | Kidney tumor tissue 1 RP | tumor tissue | kidney |
| Tissue | LD tissue sample Sus scrofa | tissue sample | longissimus dorsi |
| Tissue | Leaf tissue Brachypodium distachyon accession ABR6 | Leaf tissue | fourth and fifth leaf |
| Tissue | Leaf transcriptome Euphorbia pekinensis (replicate = biological replicate 2) | Leaf transcriptome | leaf (replicate = biological replicate 2) |
| Tissue | Leave segment region 4 (photosynthetic) collected 10pm | Leave segment | leaf segment region 4 |
| Tissue | Leave segment region 5 (photosynthetic) collected 10pm | Leave segment | leaf segment region 5 |
| Tissue | MCF-10A digested nuclei biological replicate 1 | nuclei biological | breast |
| Tissue | Model organism animal sample Dichorragia nesimachus | sample Dichorragia | thoracic muscle tissue or legs |
| Tissue | Model organism animal sample litopenaeus vannamei | sample litopenaeus | gill |
| Tissue | Multispecies transcriptome raw reads Solanum lycopersicum style tissue pollinated Solanum pennellii pollen +1 days flower opening (individual L2+P1) | style tissue | style+pollen |
| Tissue | Multispecies transcriptome raw reads Solanum pennellii style tissue pollinated Solanum lycopersicum pollen +1 days flower opening (individual P3+L3) | style tissue | style+pollen |
| Tissue | Partial transcriptome abdominal tissue female | abdominal tissue | abdomen |

| | | | |
|---|---|---|---|
| | Hypolimnas bolina | | |
| Tissue | RNAseq Ptychodera flava: head regeneration 0h-3 | regeneration 0h-3 | head amputation plane, anterior to collar |
| Tissue | RNAseq Ptychodera flava: head regeneration 96h-1 | regeneration 96h-1 | head amputation plane, anterior to collar |
| Tissue | Sample ABHDF1 RNASeq bacteria-HDFa co-culture experiments | Sample ABHDF1 | adult skin |
| Tissue | Single-cell sequencing individual murine aortal smooth muscle cells [Harmandeep_SMC_run2_09] | smooth muscle | aorta |
| Tissue | Single-cell sequencing individual murine aortal smooth muscle cells [Harmandeep_SMC_run2_86] | smooth muscle | aorta |
| Tissue | Skin sample old subjects | Skin sample | skin (epidermal suction blister samples) |
| Tissue | Sunflower species whole genome shotgun sequencing | whole genome | leaves |
| Tissue | The blood sample KPGP-00336 | sample KPGP-00336 | blood |
| Tissue | The integument Dazao 16 h head capsule slippage fourth molt | head capsule | integument |
| Tissue | Thymus tissue sample male female Indian-origin rhesus macaque | tissue sample | thymus |
| Tissue | Transcriptome analysis table grape berry cv | grape berry | baya |
| Tissue | Triticum aestivum microspore embryogenesis sample | embryogenesis sample | anther in vitro culture |
| Tissue | Tumor DNA STAT1 -/- mouse model human breast cancer sample M_CA-SSM3-SSM3 | Tumor DNA | cl |
| Tissue | Tumor DNA sample human female participant Texas Cancer Research Biobank Open Access Data Sharing BioProject | Tumor DNA | pancreas |
| Tissue | WR1913 subject 10 oropharyngeal swab | WR1913 subject | oropharynx |
| Tissue | Whole genome bisulfite sequencing control CD1A dendritic cells | Whole genome | primary cells |
| Tissue | Whole genome sequence vervet Chlorocebus tantalus animal | Whole genome | blood cell pellet |
| Tissue | copper treatment control sample shoot 01 | sample shoot | shoots |
| Tissue | copper treatment treated sample root 02 | sample root | roots |
| Tissue | copper treatment treated sample shoot 02 | sample shoot | shoots |
| Tissue | islet preparation 1 control condition | islet preparation | islets of langerhans |
| Tissue | islet preparation 3 cytokine treatment | islet preparation | islets of langerhans |
| Tissue | juvenile skin biopsy | skin biopsy | whole skin |
| Tissue | muscle tissue Tibetan 279 | muscle tissue | longissimus dorsi muscle |
| Tissue | muscle tissue Wujin 48 | muscle tissue | longissimus dorsi muscle |
| Tissue | non-injured ovaries ovarian surgery model | surgery model | ovary |
| Tissue | purified Tomato-positive PDAC cancer cells mouse 0758 LN metastasis - technical replicate1 | LN metastasis | ln met (pdac) |
| Tissue | rosette plant cold treatment | plant cold | shoot apices |
| Tissue | sample ID 19 | sample ID | hippocampus |
| Tissue | sample P53hom-22c | sample P53hom-22c | tail |
| Tissue | sample P53hom-39c | sample P53hom-39c | tail |
| Tissue | sample P53hom-48c | sample P53hom-48c | tail |
| Tissue | sample Rp048c | sample Rp048c | tail |
| Tissue | scion mature leaves removed | leaves removed | developing leaf, grafted |
| Tissue | scion mature leaves removed | leaves removed | shoot apex, grafted |
| Tissue | specimen D015 | specimen D015 | regenerating tail |
| Age | 0 days old | days old | 0 |
| Age | 10 dpf old | dpf old | 10 |
| Age | 16S rDNA based microbial profile ileostoma effluent collected subject 1 morning day 1 | morning day | 79 |
| Age | 16S rRNA based microbial profile ileostoma | afternoon day | 65 |

| | | | |
|---|---|---|---|
| | effluent collected subject 2 afternoon day 1 | | |
| Age | 2 hours post exposure | hours post | juvenile |
| Age | 2 week dark-adapted | 2 week | 7 days old |
| Age | 2 week dark-adapted | 2 week | 7 days old |
| Age | 2014 Sum - mean (22101) | 2014 Sum | 3 |
| Age | 4 hr FGF + LY294002 Rep2 | 4 hr | e13.5 |
| Age | 4 hr FGF + PD325901 Rep1 | 4 hr | e13.5 |
| Age | 5 weeks old | weeks old | 5 |
| Age | 7 week age | week age | 7 weeks |
| Age | 7 week age | week age | 7 week |
| Age | 8-oxoG challenge - 60 min | 60 min | 8 weeks |
| Age | Alamo Salt Experiment Control Rep 2 - 12 h | 12 h | 30d |
| Age | Alamo Salt Experiment Treatment Rep 1 - 24 h | 24 h | 30d |
| Age | Arabidopsis thaliana Cold Treatment 12 hours | 12 hours | 21 day |
| Age | DMD sample Replicate 2 myotube day 2 | day 2 | 1 years old |
| Age | DMD sample Replicate 3 myotube day 1 | day 1 | 2 years old |
| Age | Day 14 Of Antibiotic Treatment Human Gut Metagenome | Day 14 | 68 |
| Age | Dental lamina Pre-initiation stage rep2 | Pre-initiation stage | one year old |
| Age | ENCODE biosample ENCBS080PZG: Established 1987 peripheral blood patient T-cell acute lymphoblastic leukemia (T-ALL) obtained two months prior death | months prior | 38 year |
| Age | ENCODE biosample ENCBS835VCD: Day 13 (T13) | Day 13 | 25 year |
| Age | Embryonic transcriptome domesticated turkey aflatoxin B1 1 day exposure liver sample 2 | day exposure | day 18 embryo |
| Age | Embryonic transcriptome domesticated turkey aflatoxin B1 1 day exposure liver sample 6 | day exposure | day 18 embryo |
| Age | Embryonic transcriptome domesticated turkey aflatoxin B1 1 day exposure liver sample 7 | day exposure | day 18 embryo |
| Age | Flower inflorescence buds stage I | stage I | two years |
| Age | Hepatitis B booster immunisation study - 1066 | booster immunisation | 42 |
| Age | Hepatitis B booster immunisation study - 1070 | booster immunisation | 57 |
| Age | Hepatitis B booster immunisation study - 1070 | booster immunisation | 57 |
| Age | Hepatitis B booster immunisation study - 1776 | booster immunisation | 28 |
| Age | Injured spinal cord Trachemys scripta elegans 4 dpl N2 | 4 dpl | 1 year |
| Age | Inoculated resistant cultivar 6hr post inoculation | 6hr post | 14 days |
| Age | Inoculated susceptible cultivar 24hr post inoculation | 24hr post | 14 days |
| Age | Late flowering PHYTOCHROME C mutant sample Brachypodium distachyon | Late flowering | few months |
| Age | Library 3 MiSeq Cre normal PCR 15 cycles | 15 cycles | 21 days |
| Age | May value evaluating role t(16 | May value evaluating role t(16 | 38 year |
| Age | Methylome WT Root 21 -Pi r2 | 21 -Pi r2 | 2 weeks + 21 d |
| Age | Methylome dcl3a Root 21 -Pi r2 | 21 -Pi r2 | 2 weeks + 21 d |
| Age | Microbiota infant TB15 1 month age | month age | 1 month |
| Age | Model organism animal sample Dermacentor andersoni 2 days | 2 days | adults |
| Age | Nuclear_RNA extract Differentiated Myotubes 7 days cell culture Duchenne Muscular Dystrophy (DMD) donors Caucasian origin_Nuclear_DMD_MT-7_9813_21 | 7 days | 3 |
| Age | Nuclear_RNA extract Differentiated Myotubes 7 days cell culture Healthy donors Caucasian origin_Nuclear_CTRL_MT-7_10006_11 | 7 days | 3 |
| Age | Patient 2 Day 3 | Day 3 | 21 |

| | | | |
|---|---|---|---|
| Age | RESA-Seq - WT 8h pA r3 B1 | 8h pA | 8 |
| Age | RRGD Timepoint VI - recovery 15 min | 15 min | 23 days |
| Age | Retina Aipl1 knockout mouse postnatal day 50 | postnatal day | 50 days |
| Age | Single cell 4 month old LMPP 10 | month old | 4 months |
| Age | Skeletal muscle 8-week old mouse | 8-week old | 8 weeks |
| Age | T1 - horses intensive training stage (after slow canter phase) (March) 3 | stage (after | 3 year old |
| Age | Vaginal microbiota associated preterm delivery | preterm delivery | 35 |
| Age | Vaginal microbiota associated preterm delivery | preterm delivery | 42 |
| Age | Vaginal microbiota associated preterm delivery | preterm delivery | 30 |
| Age | Vaginal microbiota associated preterm delivery | preterm delivery | 26 |
| Age | Vaginal microbiota associated preterm delivery | preterm delivery | 19 |
| Age | Vaginal microbiota associated preterm delivery | preterm delivery | 37 |
| Age | Vaginal microbiota associated preterm delivery | preterm delivery | 35 |
| Age | Vaginal microbiota associated preterm delivery | preterm delivery | 33 |
| Age | Vaginal microbiota associated preterm delivery | preterm delivery | 33 |
| Age | Vaginal microbiota associated preterm delivery | preterm delivery | 29 |
| Age | Vaginal microbiota associated preterm delivery | preterm delivery | 35 |
| Age | Vaginal microbiota associated preterm delivery | preterm delivery | 31 |
| Age | Vaginal microbiota associated preterm delivery | preterm delivery | 29 |
| Age | Vaginal microbiota associated preterm delivery | preterm delivery | 29 |
| Age | Vaginal microbiota associated preterm delivery | preterm delivery | 27 |
| Age | Vaginal microbiota associated preterm delivery | preterm delivery | 19 |
| Age | Vaginal microbiota associated preterm delivery | preterm delivery | 20 |
| Age | Vaginal microbiota associated preterm delivery | preterm delivery | 23 |
| Age | Vaginal microbiota associated preterm delivery | preterm delivery | 32 |
| Age | Vaginal microbiota associated preterm delivery | preterm delivery | 22 |
| Age | Vaginal microbiota associated preterm delivery | preterm delivery | 27 |
| Age | Vaginal microbiota associated preterm delivery | preterm delivery | 37 |
| Age | Vaginal microbiota associated preterm delivery | preterm delivery | 27 |
| Age | Vaginal microbiota associated preterm delivery | preterm delivery | 25 |
| Age | Vaginal microbiota associated preterm delivery | preterm delivery | 33 |
| Age | Vaginal microbiota associated preterm delivery | preterm delivery | 38 |
| Age | Vaginal microbiota associated preterm delivery | preterm delivery | 28 |
| Age | Vaginal microbiota associated preterm delivery | preterm delivery | 33 |
| Age | Vaginal microbiota associated preterm delivery | preterm delivery | 24 |
| Age | Vaginal microbiota associated preterm delivery | preterm delivery | 33 |
| Age | Vaginal microbiota associated preterm delivery | preterm delivery | 17 |
| Age | Vaginal microbiota associated preterm delivery | preterm delivery | 20 |
| Age | Vaginal microbiota associated preterm delivery | preterm delivery | 21 |
| Age | Vaginal microbiota associated preterm delivery | preterm delivery | 36 |
| Age | Vaginal microbiota associated preterm delivery | preterm delivery | 24 |
| Age | Vaginal microbiota associated preterm delivery | preterm delivery | 28 |
| Age | Vaginal microbiota associated preterm delivery | preterm delivery | 22 |
| Age | Vaginal microbiota associated preterm delivery | preterm delivery | 23 |
| Age | Vaginal microbiota associated preterm delivery | preterm delivery | 33 |
| Age | Vaginal microbiota associated preterm delivery | preterm delivery | 38 |
| Age | Vaginal microbiota associated preterm delivery | preterm delivery | 26 |
| Age | Vaginal microbiota associated preterm delivery | preterm delivery | 34 |
| Age | Vaginal microbiota associated preterm delivery | preterm delivery | 35 |
| Age | Vaginal microbiota associated preterm delivery | preterm delivery | 37 |
| Age | Vaginal microbiota associated preterm delivery | preterm delivery | 27 |
| Age | Vaginal microbiota associated preterm delivery | preterm delivery | 29 |
| Age | Vaginal microbiota associated preterm delivery | preterm delivery | 24 |
| Age | Vaginal microbiota associated preterm delivery | preterm delivery | 31 |
| Age | Vaginal microbiota associated preterm delivery | preterm delivery | 37 |
| Age | Vaginal microbiota associated preterm delivery | preterm delivery | 25 |

| | | | |
|---|---|---|---|
| Age | Vaginal microbiota associated preterm delivery | preterm delivery | 25 |
| Age | Vaginal microbiota associated preterm delivery | preterm delivery | 29 |
| Age | Vaginal microbiota associated preterm delivery | preterm delivery | 22 |
| Age | Vaginal microbiota associated preterm delivery | preterm delivery | 22 |
| Age | Vaginal microbiota associated preterm delivery | preterm delivery | 22 |
| Age | Vaginal microbiota associated preterm delivery | preterm delivery | 38 |
| Age | Vaginal microbiota associated preterm delivery | preterm delivery | 42 |
| Age | Vaginal microbiota associated preterm delivery | preterm delivery | 35 |
| Age | Vaginal microbiota associated preterm delivery | preterm delivery | 24 |
| Age | Vaginal microbiota associated preterm delivery | preterm delivery | 31 |
| Age | Vaginal microbiota associated preterm delivery | preterm delivery | 21 |
| Age | Zeitgeber Time 00 | Zeitgeber Time | 3 weeks |
| Age | Zeitgeber Time 18 | Zeitgeber Time | 5 weeks |
| Age | bisulfite treated genomic DNA old sun exposed epidermis 3 | old sun | 83 |
| Age | cinerea B9 different inoculation time treatments | time treatments | 1 year |
| Age | five 5w old individuals (male) | 5w old | 5 week |
| Age | time point 2 | time point | 65.397 |
| Age | zebrafish normal developmental age 72hpf control rep2 | developmental age | 72 hpf¬£¬®hour post fertilization¬£¬© |
| Data type | Non-tumor DNA sample human male participant dbGaP study "Whole Exome Sequencing Chronic Lymphocytic Leukemia" | Exome Sequencing Chronic | snv aggregate (.maf) |
| Data type | Tumor DNA sample human male participant dbGaP study "Whole Exome Sequencing Chronic Lymphocytic Leukemia" | Exome Sequencing Chronic | snv aggregate (.maf) |
| Platform | 16S rDNA amplicon patient 11 | 16S rDNA amplicon | 454 |
| Platform | 16S rDNA amplicon patient 5 | 16S rDNA amplicon | 454 |
| Platform | 16S rRNA control root 3 | 16S rRNA | ion torrent |
| Platform | 16S rRNA mycorrhiza leaf 1 | 16S rRNA | ion torrent |
| Platform | 16S rRNA sequences fecal bacterial community Chicken | 16S rRNA | 454 gs flx |
| Platform | Mouse fecal sample VLP-purification 454 Sequencing | 454 Sequencing | 454 flx titanium |
| Platform | Mouse fecal sample VLP-purification 454 Sequencing | 454 Sequencing | 454 flx titanium |
| Platform | Pyrosequencing 16S rRNA amplicons pig gastrointestinal mucosal scrapings | Pyrosequencing 16S rRNA amplicons | pyrosequncing |
| Platform | Pyrosequencing 16S rRNA amplicons pig gastrointestinal mucosal scrapings | Pyrosequencing 16S rRNA amplicons | pyrosequncing |
| Platform | Pyrosequencing 16S rRNA amplicons pig gastrointestinal mucosal scrapings | Pyrosequencing 16S rRNA amplicons | pyrosequncing |
| Platform | SAG SCGC AB-704-M06 partial SSU rRNA gene | SSU rRNA | sanger |
| Platform | SAG SCGC AB-704-N13 partial SSU rRNA gene | SSU rRNA | sanger |
| Platform | SAG SCGC AB-706-C03 partial SSU rRNA gene | SSU rRNA | sanger |
| Platform | SAG SCGC AB-706-E02 partial SSU rRNA gene | SSU rRNA | sanger |
| Platform | SAG SCGC AB-706-E13 partial SSU rRNA gene | SSU rRNA | sanger |
| Platform | SAG SCGC AB-706-E22 partial SSU rRNA gene | SSU rRNA | sanger |
| Platform | SAG SCGC AB-706-F21 partial SSU rRNA gene | SSU rRNA | sanger |
| Platform | SAG SCGC AB-706-K06 partial SSU rRNA gene | SSU rRNA | sanger |
| Platform | SAG SCGC AB-706-M02 partial SSU rRNA gene | SSU rRNA | sanger |
